# *In silico* design of Multi-epitope-based peptide vaccine against SARS-CoV-2 using its spike protein

**DOI:** 10.1101/2020.04.23.055467

**Authors:** Debarghya Mitra, Janmejay Pandey, Alok Jain, Shiv Swaroop

## Abstract

SARS-CoV-2 has been efficient in ensuring that many countries are brought to a standstill. With repercussions ranging from rampant mortality, fear, paranoia, and economic recession, the virus has brought together countries to look at possible therapeutic countermeasures. With prophylactic interventions possibly months away from being particularly effective, a slew of measures and possibilities concerning the design of vaccines are being worked upon. We attempted a structure-based approach utilizing a combination of epitope prediction servers and Molecular dynamic (MD) simulations to develop a multi-epitope-based subunit vaccine that involves the two subunits of the spike glycoprotein of SARS-CoV-2 (S1 and S2) coupled with a substantially effective chimeric adjuvant to create stable vaccine constructs. The designed constructs were evaluated based on their docking with Toll-Like Receptor (TLR) 4. Our findings provide an epitope-based peptide fragment that can be a potential candidate for the development of a vaccine against SARS-CoV-2. Recent experimental studies based on determining immunodominant regions across the spike glycoprotein of SARS-CoV-2 indicate the presence of the predicted epitopes included in this study.

## 1. Introduction

In December 2019, the town of Wuhan in Hubei, China, experienced an outbreak of a highly pathogenic virus prevalent with high transmissibility across humans and potentially responsible for varied symptoms associated with respiratory-based complications. Months after the outbreak, the disease has spread across the globe, infecting 14 million people and responsible for the deaths of another 5.9 lacs (at submission). It has all been implicated in a pathogen belonging to the genus *Betacoronavirus* of the Coronaviridae family and has been identified as the severe acute respiratory syndrome coronavirus-2 (SARS-CoV-2). The WHO declared it a pandemic on 12^th^ March, 2020 allowing countries across the world to take appropriate measures to fend themselves from the disease and prevent its spread.

The disease transmission has left very few countries free from the repercussions of the disease with surmounting pressure on hospital settings and resources to contain this public health emergency. The scientific community is on an overdrive to look into prophylactic measures as the first line of defense against the disease. The development of rapid diagnostics to determine and isolate disease affected individuals along with repurposing and identifying existing FDA-approved drugs to provide substantial treatment are underway. Still, going by the looks of it and the unprecedented pace of development of therapeutics, it would require several months to cross over several clinical trials and finally be made available to the population.

The study of SARS-CoV-2 has been easier due to the manifestation of a similar homolog of the disease SARS-CoV more than a decade ago and comprises of similar structural counterparts that aid in the process of infection, including a spike glycoprotein, membrane glycoprotein, an envelope protein and a nucleocapsid protein[1, 2]. The genome sequence and its analysis were done promptly and made available [3, 4]. Several structures over SARS-Cov-2 S protein in interaction with human ACE-2 (hACE-2) have been made publicly available over the RCSB PDB[5–8], allowing the scientific community to proceed towards the development of probable therapeutic measures ranging from repurposing drugs[9], design of small molecule inhibitors, identification of unique targets and elucidating molecular mechanisms of the viral protein machinery[10] and how it establishes itself inside the host body. The interaction between the spike glycoprotein and the human ACE-2 receptor is important as it initiates the process of viral entry into human beings through contact with an infected individual. The identification of a probable site upstream of the receptor-binding motif (which includes the residues on S1 that interact with hACE-2), which undergoes cleavage and requires priming by TMPRSS2 before stable fusion of the viral membrane complex to the hACE2[11] and involves sufficient nonreversible conformational changes that put into motion the process of viral entry[1, 5].

A vaccine can be a useful measure against this positive, single-stranded RNA virus. Several candidate vaccines are already underway since they are considered the only reasonable step in immunizing individuals worldwide and stopping the disease from steamrolling through countries. This paper investigates the utilization of the spike glycoprotein of SARS-CoV-2 as a potential immunogen for designing a multi-epitope peptide-based subunit vaccine with a chimeric adjuvant in tow. The spike glycoprotein is evidently an appropriate immunogen capable of eliciting neutralizing antibodies, which will inadvertently lead to the establishment and elicit an immune response to prevent viral entry and act as a prophylactic [6].

The work envisages the prediction of significant epitopes and the design of a multi-epitope-based peptide subunit vaccine coupled with a chimeric adjuvant. The immunogen considered involves the two domains of the spike glycoprotein involved in binding with hACE-2 followed by viral and cellular membrane fusion through S1 and S2, respectively. The cleavage site present between S1 and S2 post-fusion has not been considered due to previous apprehensions on the immune system’s hypersensitivity with reference to the entire spike glycoprotein of SARS-CoV. The utilization of these epitopes and the vaccine construct across an experimental setting will help provide essential evaluation and aid in developing a vaccine to generate a robust immunological prophylactic response against SARS-CoV-2.

## 2. Methods

### 2.1 Sequence Retrieval for SARS-CoV-2 Spike Glycoprotein

The SARS-CoV-2 spike glycoprotein sequence (PDB ID: 6VSB)[6] was checked for specific subunits and domains. S1 subunit (27-526 residues) comprising the Receptor Binding Domain (331-524 residues) along with the S2 subunit (663-1146) were identified from Wrapp et al[6]. The corresponding structure submitted over RCSB PDB[6] was utilized for visualization of the SARS-CoV-2 spike protein using UCSF Chimera [12].

### 2.2 Prediction of B-cell epitopes

The sequences of S1 and S2 domains were separately used to determine linear B cell epitopes through different servers. We used the Artificial Neural Network (ANN) approach-based server ABCPred[13], Bcepred[14], and the Immune Epitope Database and Analysis Resource (IEDB) based linear B cell epitope prediction tools[15–20] to predict probable epitope sequences across the query sequences. The latter two servers utilize physicochemical parameters like hydrophilicity, polarity, surface accessibility, and flexibility to predict a B cell epitope as has been previously evidenced. The predicted linear B cell sequences from ABCPred were matched with the probable epitopes predicted over the IEDB-based Bepipred 2.0 based linear prediction of epitopes for the different parameters inclusive of surface accessibility area, hydrophilicity, polarity and flexibility, and the results from the two servers were corroborated. These same parameters were then again checked with the predicted epitopes by Bcepred. Consensus sequences that emerged from all three were considered to be the potential epitopes in this regard. Since conformational or discontinuous B cell epitopes could be predicted through ElliPro [21] and Discotope [22], they were utilized for analysis of the consensus sequences arrived upon earlier. Consensus 16-mer epitopes were predicted using these servers.

### 2.3 Prediction of MHC II based Helper T Lymphocytes (CD4^+^)

The probable MHC II binding epitopes on S1 and S2 domains were predicted, and a consensus approach was utilized again to arrive at the conclusive sequences. The prediction of helper T lymphocytes was made using MHCPred[23], SYFPEITHI[24], NetMHCIIpan 3.2 server [25], and the IEDB server [26, 27]. A consensus selection of epitopes from the four different servers allowed us to improve upon and circumvent the limitations associated with the prediction of MHC II binders due to their polymorphic nature, peptide length variation, and determination of the appropriate peptide-binding core. Hence, looking into the limitations, these servers were determined to be the best amongst available servers that can be utilized in this study. Each of these prediction servers was compared with experimental datasets to assess their performance. SYFPEITHI predicts nonamer sequences based on the weighted contribution of each amino acid sequence present across a predicted epitope sequence. MHCPred allows the prediction of 9-mer epitopes based on multivariate statistical methods, and NetMHCIIpan 3.2 enables the determination of 15-mer epitope fragments but with limited allele-specific choices. The IEDB-based MHC II binding epitope sequence prediction server was also utilized because it employs a combination of methods ranging from ANN to SMM and ranks probable epitopes on the basis of a percentile score and an IC_50_ score. The sequences predicted over the different servers were either 15 mers or 9 mers depending on the server used. Based on the consensus selection across these platforms and overlapping regions of the predicted epitope sequences, uniform 15-mer epitopes were selected.

### 2.4 Prediction of MHC I based Cytotoxic T Lymphocytes (CD8^+^)

The prediction of Cytotoxic T Lymphocyte cells (T_CTL_) through MHC I binding servers involved the utilization of NetMHC 4.0 [28], MHC-NP[29], NetCTL 1.2[30], and the IEDB-based T cell epitope tools[31]. All these servers predict a nonamer epitope sequence using a default dataset along with probable interacting human leukocyte antigen alleles with SARS-CoV identified from the literature. Employing a consensus selection of the predictions from the four servers, we were able to list the appropriate T_CTL_ epitopes with relative confidence. The NetMHC 4.0 utilizes an Artificial Neural Network to predict epitopes, NetCTL 1.2 server allows the identification of epitopes based on improved datasets with a sensitivity of 0.80 and specificity of 0.97 across the filtering threshold employed, MHC-NP employs a Machine Learning approach towards the prediction of naturally processed epitopes by the MHC, whereas IEDB-based prediction of T cell epitopes sorts epitopes based on the percentile score and low IC_50_ values across a combination of ANN and SMM approaches based on an appropriate peptide library. The MHC I alleles specific for a SARS-CoV-2 manifestation were based on the alleles confirmed during the outbreak of SARS-CoV at the beginning of the millennium.

### 2.5 Validation of predicted epitopes

The consensus sequences thus arrived upon can be considered as being capable of eliciting the necessary immune response. Additionally, each of the sequences was filtered based on their predicted antigenicity over Vaxijen[32], allergenicity over AllerTop[33] and Algpred[34], toxicity over ToxinPred[35], ability of eliciting Interferon-gamma over IFN-G[36] servers. Also, the epitope sequences were matched with human proteins to prevent cases of antibody response against a self-antigen with the Multiple Peptide Match Tool. Each of the epitope sequences was matched with their corresponding secondary structure over the cryoEM structure (PDB ID: 6VSB)[6] and listed as either alpha-helices, coils, strands or beta-sheets. This was done to allow for sequence arrangement for improved molecular modeling. Based on the conservation of the surface glycoprotein across the different strains deposited over the GISAID repository (https://www.gisaid.org/about-us/mission/) and the opinion that not much variability has been observed across these strains, we checked the coverage of the epitope sequences we have predicted with the sequences from the repository of surface glycoprotein and also the genome sequence deposited over NCBI from India [37].

### 2.6 Design of Adjuvant

In a peptide-based subunit vaccine, the predicted B and T cell epitopes are not sufficient to elicit a strong immune response to generate the necessary prophylactic measures. Therefore, a suitable adjuvant must also be added to the vaccine design[38]. Recently, several approaches have been made to utilize a chimeric adjuvant comprising two or more separately identified adjuvants [39, 40]. We used a similar approach and carried out an *in silico* design comprising of three different adjuvants that have been used earlier in different vaccines or as agonists of TLR-4 receptors. A recent review of suitable adjuvants for different TLRs using this approach has been published[41]. It was backed up by experimental evidence of the identified adjuvants capable of eliciting an immune response on interaction with toll-like receptors (TLRs) separately and downstream immune signaling generating the necessary prophylactic measures that can be expected from a vaccine against SARS-CoV-2 [42]. The generation of neutralization antibodies by the peptide sequences of the surface glycoprotein has already been evidenced in previous instances. Hence, their linking with a triagonist chimeric adjuvant supports their utilization as a suitable prophylactic measure. A suitable rigid linker was utilized to join the peptide sequence and their position was rearranged at the N-terminal to design several constructs of the vaccine. A single construct with adjuvants at the N and C-terminal of the vaccine was also considered.

### 2.7 Design and characterization of Vaccine Constructs

Appropriate linkers were utilized to join intra B cell epitopes, T-Helper, and T-Cytotoxic epitopes and also between them in the vaccine construct[43, 44]. The designed construct comprises the B cell epitopes linked to the adjuvant through a rigid EAAAK linker (helix forming) toward the N-terminal, followed by a GPGPG linker connecting the T_HTL_ epitopes and an AAY linker for the T_CTL_ epitopes toward the C-terminal. A DP linker was utilized to link the three adjuvants at the N terminal. The three adjuvants were rearranged each time, and the vaccine constructs were analyzed for physicochemical properties using ProtPARAM tool[45], allergenicity[33], antigenicity[32], and scanned for probable stimulants of interferon gamma[36]. The ToxinPred database[35] was used to analyze each of the units of the vaccine construct and hence each of these constructs can be considered nontoxic.

### 2.8 Molecular Modeling of Vaccine Constructs

The main challenge associated with the molecular modeling of the vaccine constructs was that the adjuvant-specific region and the epitope-specific region matched with two different templates. Although in both cases there was sufficient sequence coverage that calls for homology modeling of the vaccine constructs, the multi-template alignment led to a modeled structure comprising mainly of strands, which would have made the modeled constructs unstable. An initial approach included modeling the two parts of the vaccine construct separately using a single template, but linking them through loops and energy minimizations of these constructs fell through. The ROBETTA server[46], which allows for comparative modeling, brought the most promising model of the vaccine constructs with sufficient secondary structure coverage. Each of these constructs was then deposited over the GALAXY web server[47] for refinement. Based on select parameters on which each modeled construct was refined, the best structures were then carried forward for validation through Ramachandran Plot, Z-score over PROCHECK[48] and ProSA-web server[49]. Additional validation was carried out using ERRAT-3D server[50]. Each of the modeled vaccine constructs was also checked for sequence mismatches.

### 2.9 Stability of the Modeled Vaccine Constructs

Each of these constructs was then assessed for its stability using the ProtParam server. The various markers for stability are made available through the Instability index (II) [51], PEST hypothesis[52], *in vivo* half-life, and the isoelectric point[53]. The assessments based on Ramachandran Plot, ERRAT 3D[50], WHATCHECK[54], and the ProSA-web server[49] allowed us to select a single vaccine construct for a prolonged molecular dynamic simulation to determine the stability of the construct and possible use in *in vivo* settings.

### 2.10 Molecular dynamics simulations

MD simulations were performed to obtain elaborate insights on the dynamic stability of the constructs using the GROMACS suits and GROMOS 54A7 force field[55, 56]. Constructs were subjected to minimization in a cubical box with the SPC/E water model using the steepest descent followed by the conjugate gradient algorithm[56]. Counter-ions were added to neutralize the system. Temperature and pressure were maintained using the modified Berendsen thermostat and Parrinello-Rahman barostat at 300K and 1 bar, respectively [57, 58]. All the systems were run for 100ns in the NPT ensemble before short equilibration of 2 ns in NVT and NPT ensembles. Long-range electrostatic interactions were evaluated using Particle Mesh Ewald (PME) algorithm with a cutoff of 1.4nm, PME order 4 and fourier spacing 0.16 [59]. Short-range interactions were evaluated up to 1.4nm. Water hydrogens and other bonds were constrained by employing SETTLE and LINCS algorithm, respectively[60, 61]. Periodic boundary conditions were applied in all three (X, Y and Z) directions. Trajectories were analyzed and visualized using GROMACS inbuilt tools and VMD[62].

### 2.11 Molecular Docking of Vaccine Construct with TLR-4

The rationale behind selecting TLR-4 (PDB ID: 3FXI_A)[63] is the fact that in the case of SARS-CoV, HIV, Influenzae, and other RNA-based viruses, this Toll-like receptor has been experimentally evidenced to be implicated[64]. TLR-4 structure was prepared for docking by removing all water molecules and bound ligands, adding polar hydrogen atoms and optimization at physiological pH 7.4 on UCSF Chimera. No other methods were employed on the TLR-4 structure prior to docking. The binding sites over TLR-4 were determined through available PDB IDs that showed binding of an adjuvant lipopolysaccharide with the Toll-like receptor and also through ElliPro[21] and Castp[65]. It was carried out through Patchdock[66] and Firedock[67].

### 2.12 MM/GBSA based evaluation of Docked Pose

Post docking, the top pose was evaluated through Molecular Mechanics/Generalized Born Surface Area (MM/GBSA) calculations over the HawkDock Server[68]. The server allows the docked pose to be assessed on a per residue basis across Van der Waal potentials, electrostatic potentials, polar solvation free-energy, and solvation free energy through empirical models. The docked pose is minimized for 5000 steps, including 2000 cycles of steepest descents and 3000 cycles of conjugate gradient minimizations based on an implicit solvent model, the ff02 force field.

## 3. Results

### 3.1 Sequence Retrieval for SARS-CoV-2 Spike Glycoprotein

The SARS-CoV-2 surface glycoprotein sequence was retrieved from PDB ID: 6VSB in FASTA format. The UCSF Chimera visualization software[12] was utilized to edit the structure and include the S1 and the S2 domains as separate entities. The use of S1 and S2 domains separately without including the cleavage site, which falls in between the two domains **(Figure 1)** for the development of an immunogen is due to previously evidenced cases of increased immune hypersensitivity utilizing the entire length of spike glycoprotein as a vaccine immunogen. [69]

**Figure 1:**
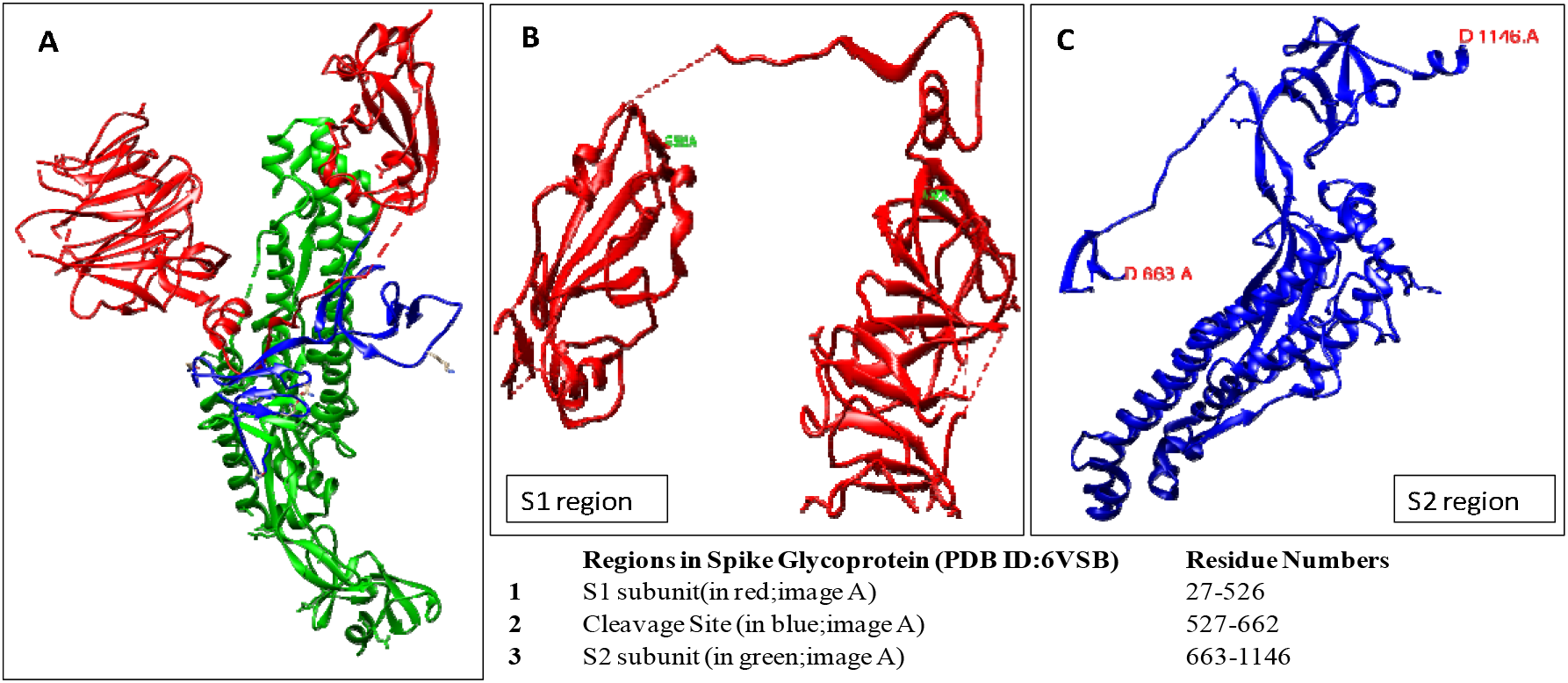
The S1 & S2 domains used in the investigation are reproduced from PDB ID: 6VSB.

### 3.2 Prediction of B cell, T_HTL_, and T_CTL_ Epitopes

The selected epitopes were based on multiple servers which used different algorithms that lead each server to arrive at the predicted epitope. Based on the stringent filters applied in each case, the epitopes predicted over the different servers were manually curated to reach upon the consensus sequences in each case. All the epitope sequences specific for S1 and S2, separately predicted by the servers, are listed. In the case of B cell [13, 14, 16–20] and MHC II [23–27] epitopes, 15 mers epitope sequences were generated, while in the case of MHC I [28–31] epitope sequences, nonamer fragments were generated.

The predicted sequences were matched with the data generated by IEDB [70] on significant epitopes present on the surface glycoprotein, which did mention the need for analysis of the predicted sites and hence the utility of the stringent filters over the different servers followed by validation. The utilization of specific alleles that have been identified in the previous manifestation of the SARS-CoV has helped us sort the predicted T cell epitopes appropriately on the different servers, which otherwise would have made prediction biased [71–74]. The selected epitopes are listed in **Table 1**.

### 3.3 Validation and Filtering of selected epitopes

The various parameters against which the selected epitopes and adjuvants were filtered included antigenicity, allergenicity, toxicity, the capability of generating interferon-gamma response (**Tables 1A, B, and C**), and chances of being identified as self-antigen by the human immune system. Among the epitope sequences filtered, 100 percent sequence coverage was detected in all but single residue mutations in 4 of them. These residues were not considered in the vaccine constructs, and the aberrations were generally isolated to the S1 domain only (Figure 2B). Hence, sequence coverage based validation of the selected epitopes was also carried out. The list of epitopes considered for the design of the vaccines did not include any sequences that did not conform to the 100% sequence coverage. The percent identity matrix was utilized to indicate that the Indian sequence shows more than 90% identity with the sequence retrieved from PDB ID:6VSB, and the epitopes utilized show 100% sequence coverage across the sequence submitted from India of the spike glycoproteins[37] **(Figure 2)**.

**Table 1(A):** Predicted and Selected B cell Epitopes from S1 and S2

| B cell Epitopes |  |  |  |  |  |  |  |
| --- | --- | --- | --- | --- | --- | --- | --- |
| Sequence | Antigen | Allergen | IFN-G | Toxin | Mol Wt (Dalton) | pI | Length (amino acids) |
| YVGYLQPRTFLLKYNE | positive | Negative | positive | negative | 2004.32 | 8.43 | 16 |
| EVRQIAPGQTGKIADY | positive | Negative | positive | negative | 1745.95 | 6.17 | 16 |
| TGKIADYNYKLPDDFT | positive | Negative | positive | negative | 1861.04 | 4.43 | 16 |
| YFKIYSKHTPINLVRD | positive | Negative | positive | negative | 1994.32 | 9.53 | 16 |
| TVEKGIYQTSNFRVQP | positive | Negative | positive | negative | 1867.09 | 8.26 | 16 |
| HVTYVPAQEKNFTTAP | positive | Negative | positive | negative | 1803 | 6.75 | 16 |

**Table 1(B):**
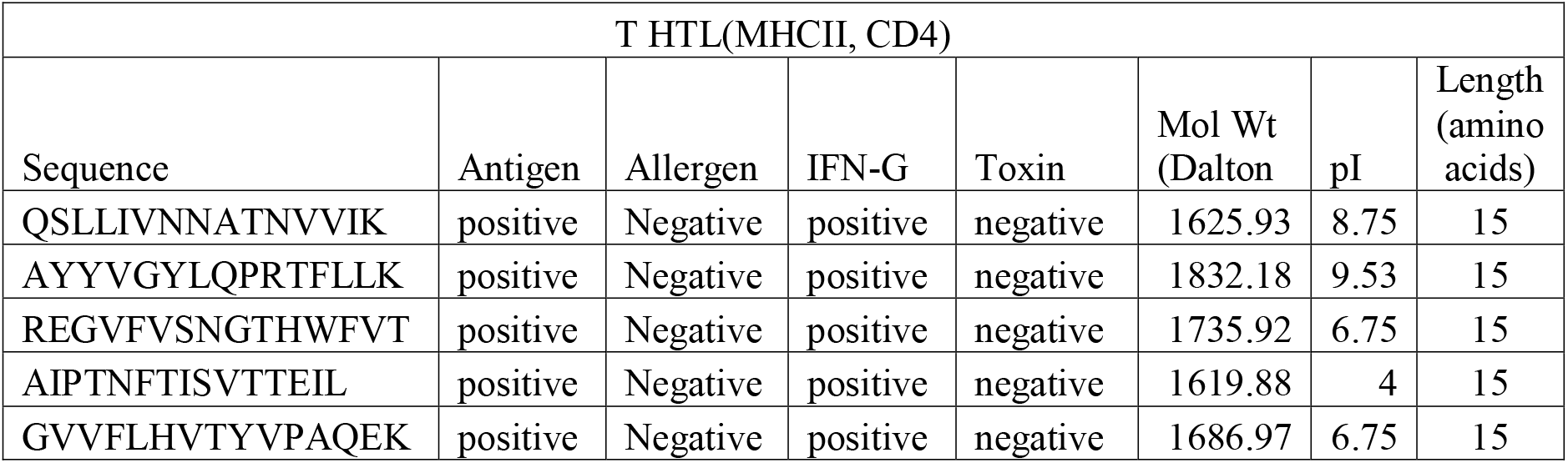
Predicted and Selected T_HTL_ cell Epitopes from S1 and S2

**Table 1(C):**
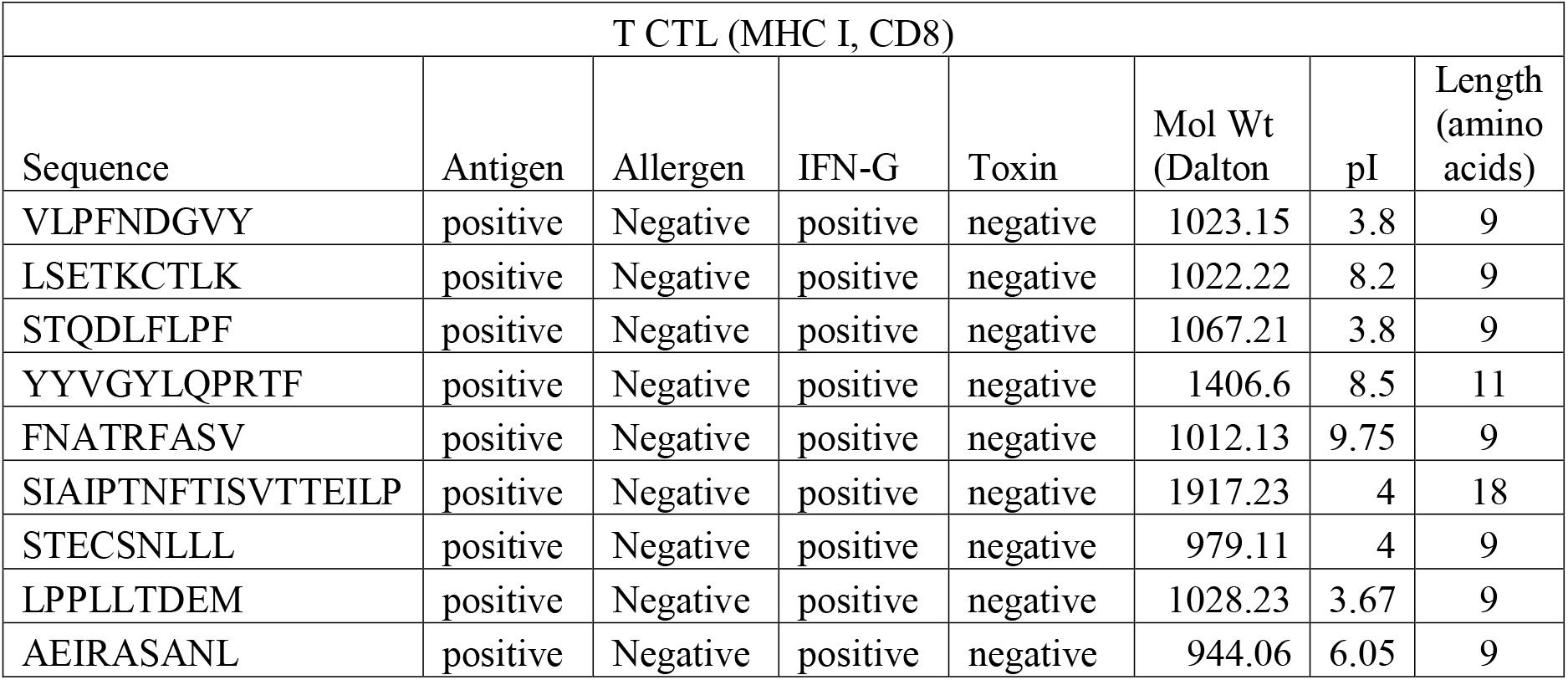
Predicted and Selected T_CTL_ cell Epitopes from S1 and S2

**Figure 2:**
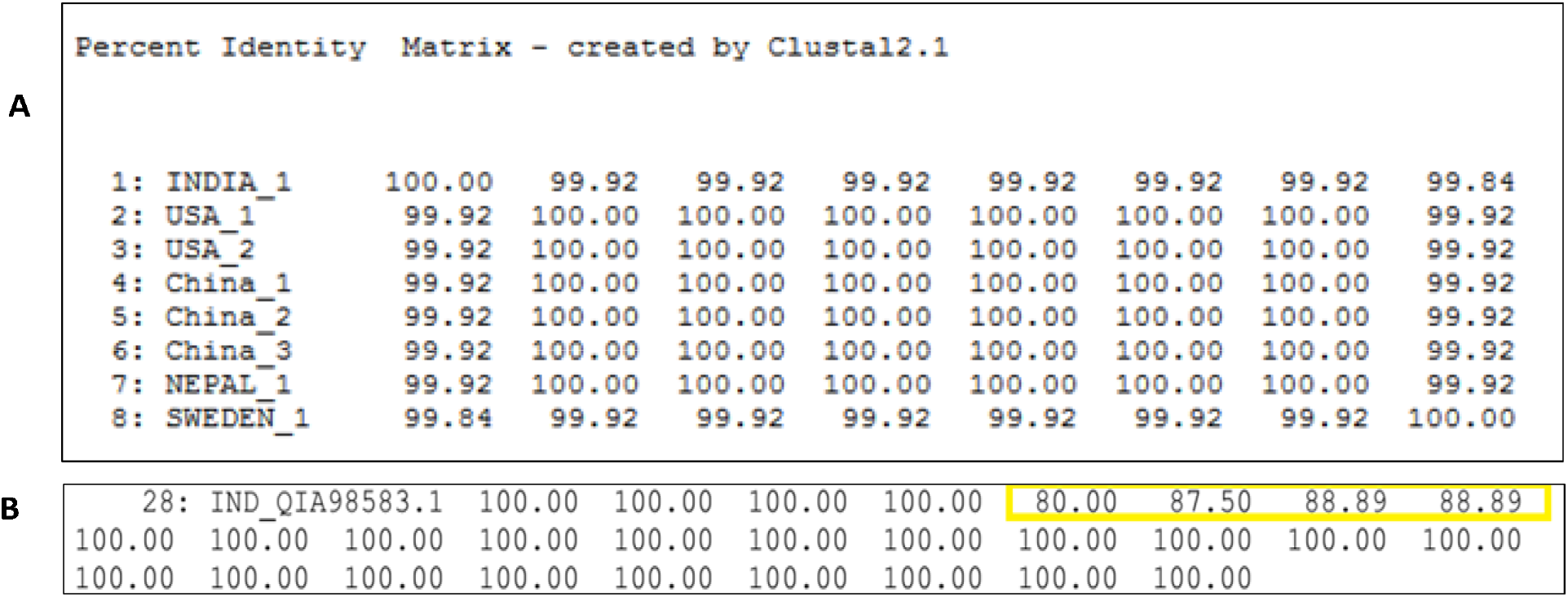
Percent Identity Matrix (PIM) of SARS-CoV-2 spike glycoprotein from GISAID (A) and (B) represents that apart from four of the epitopes selected, all have 100% sequence coverage across the submitted Indian sequence of SARS-CoV-2. The overall sequence similarity of the submitted Indian sequence across the 7 other sequences from GISAID indicates the overall coverage sequence of the epitopes selected for vaccine design falls within acceptable parameters.

### 3.4 Design of Adjuvant

The triagonist chimeric adjuvant has the propensity to elicit immune responses to allow for its use as a suitable adjuvant against SARS-CoV-2. RS09[75], TR-433[76], and an Mtb based Hsp70 partial sequence[77, 78] were utilized in this endeavor. An appropriate rigid polar linker (DP) was utilized in this regard. Among the designed constructs, all of the rearrangements were validated based on their antigenicity score predicted by Vaxijen[32]. The adjuvants considered were all verified as TLR-4 agonists. The use of fragments of *Mycobacterial* hsp70 towards the generation of cytokines and natural killer cells, and also for antigen-specific CTL responses has been verified [79–81]. The initial constructs and the parameters across which they were assessed are mentioned in **Supplementary Table 1A and 1B**. The use of lipopolysaccharide based adjuvants has been abrogated in this study due to constraints of utilizing modeling, docking, and simulation-based studies from their perspective.

### 3.5 Design and Characterization of Vaccine Constructs

The arrangement and validation of antigenicity of the vaccine constructs based on the linkers and the adjuvants led to the terminal selection of 5 different vaccine constructs. These constructs were then subsequently characterized based on the ProtParam parameters, which determined the molecular weight and isoelectric point and predicted each to be a stable construct. Also, the constructs were found to be suitably antigenic, nontoxic, and non-allergens. Regions of the vaccine constructs were predicted to have sufficient B cell epitopes and capable of generating interferon-gamma response, and have 100% sequence coverage **(Table 2 and Figure 3)**.

**Table 2:**
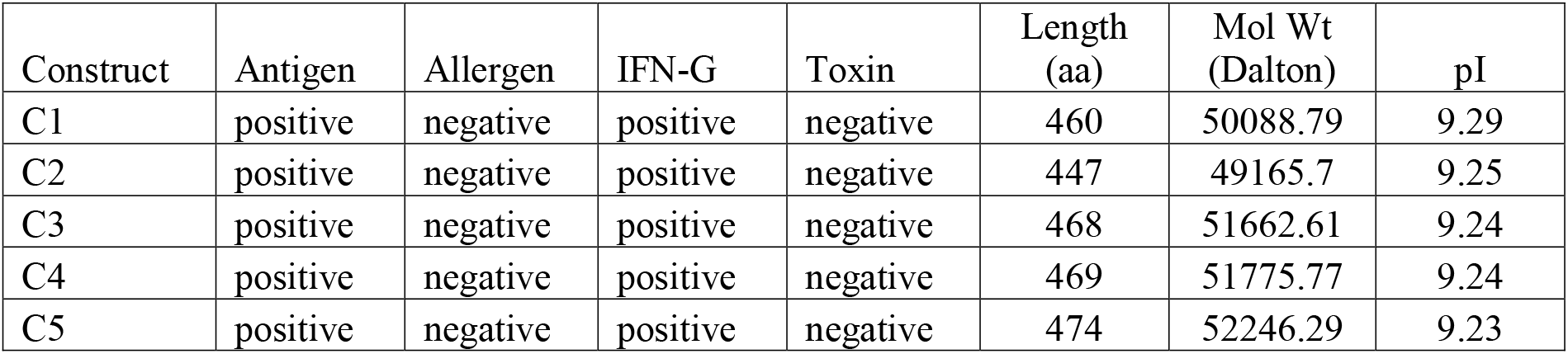
Validation of the five major vaccine constructs designed

**Figure 3:**
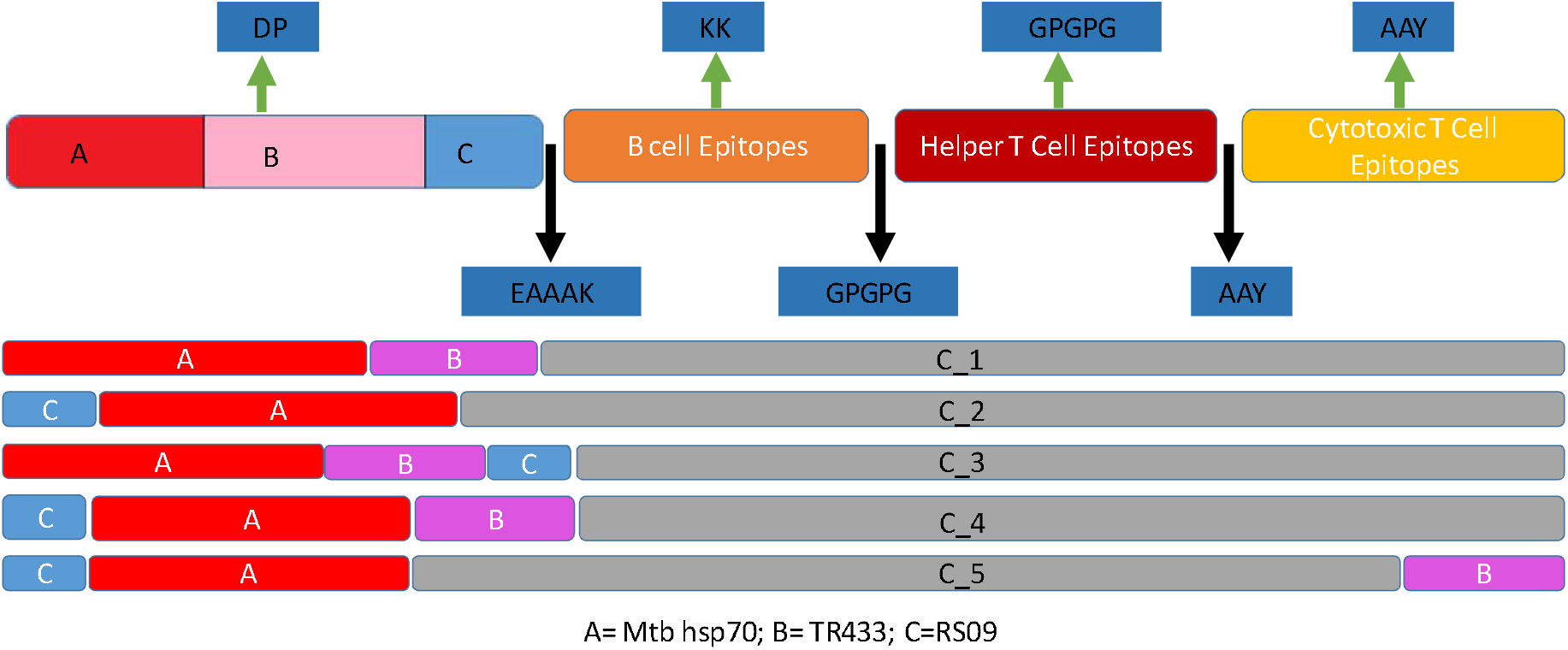
Designed vaccine constructs from C_1 to C_5 with linkers and adjuvants. Vaccine construct arrangement A, B, and C refer to the three different adjuvants used. Designed Constructs based on rearrangement and numbered appropriately.

### 3.6 Molecular Modeling and Validation of Vaccine Constructs

The vaccine constructs thus designed were made through comparative modeling over ROBETTA, and revealed a structure better than the other platforms utilized in this regard **Figure 4**.

**Figure 4:**
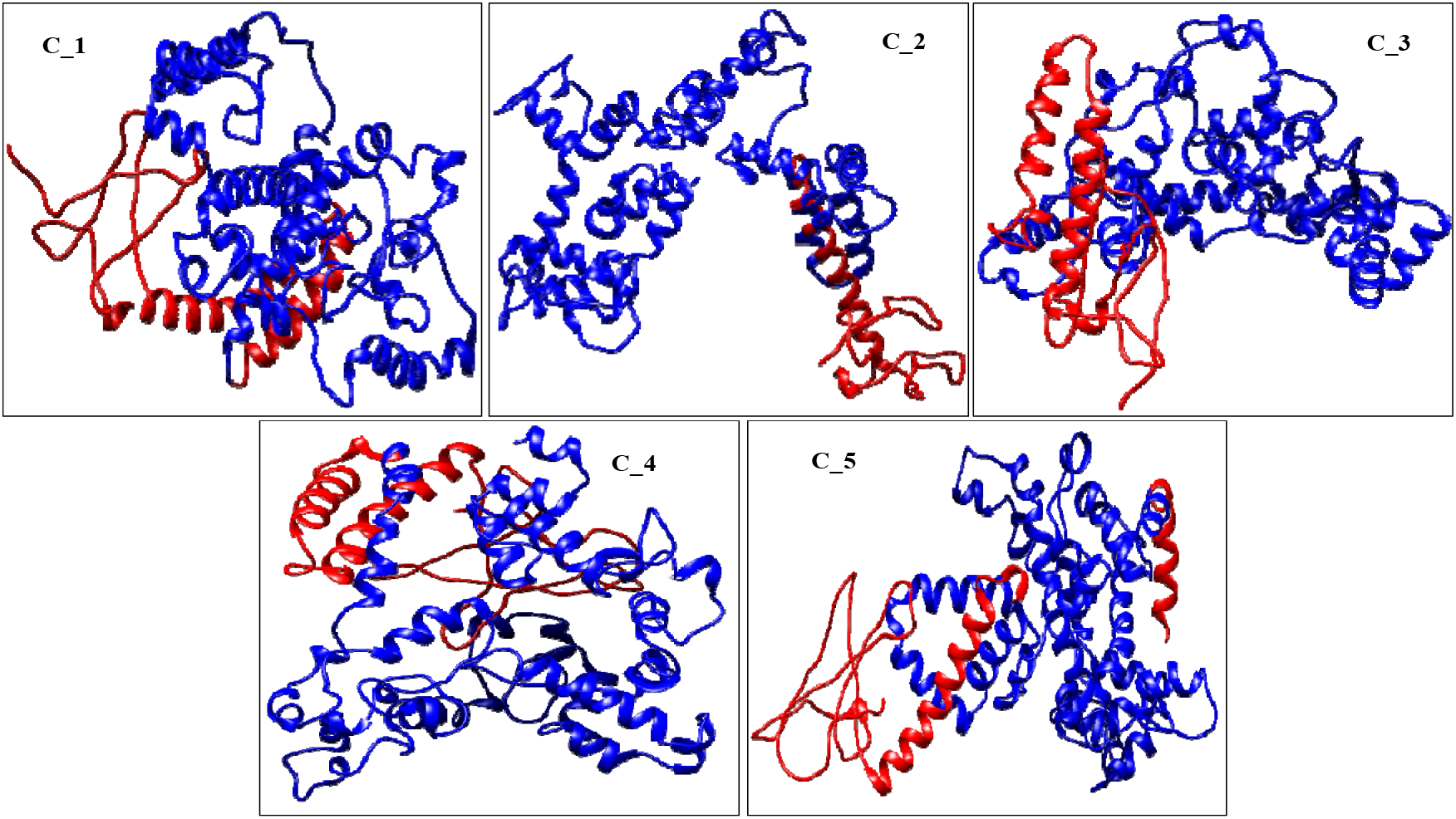
CM based modeling of designed vaccine constructs over ROBETTA server. The blue regions are the epitope sequences with the red indicating the adjuvants used and their positions.

Validation and refinement of the modeled structures revealed a stable construct in each case, which can be prominently detected by the predominant presence of defined secondary structures across the span of the construct interspersed by coils and helices. The modeled constructs were refined through the GALAXY server **(Table 3)** and validated through Ramachandran Plots and Z-score from ProSA-webserver **(Table 4C)**.

**Table 3:**
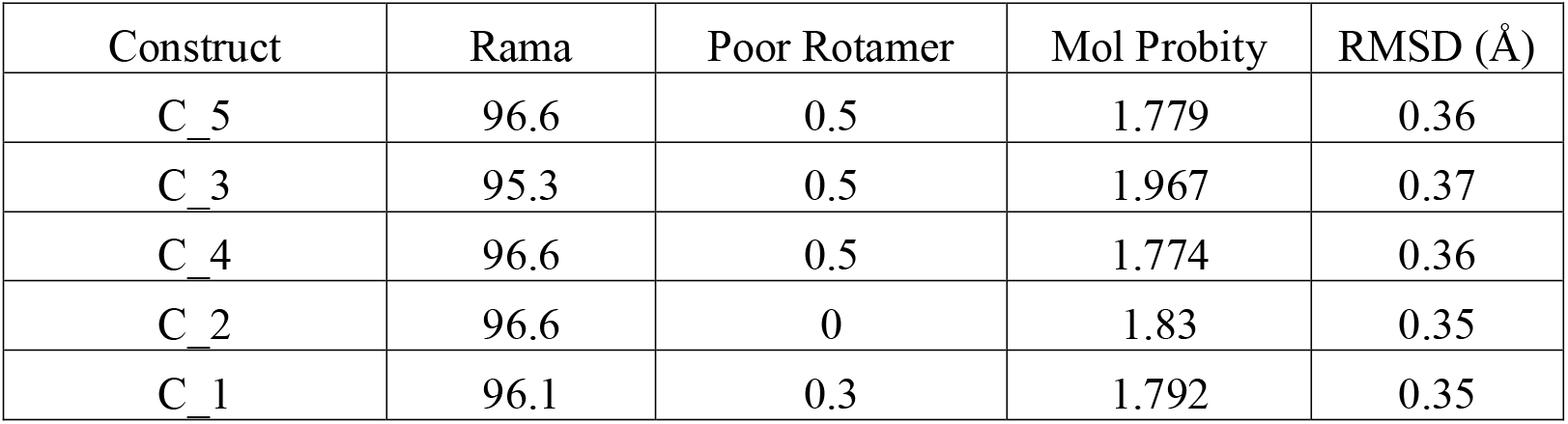
Refinement based on GALAXY SERVER of modeled constructs

### 3.7 Stability of the Modeled Vaccine Constructs

Attempts were made to assess the stability of the designed constructs before moving on to docking and simulation studies with TLRs. Since it comprises three different adjuvants, an ensemble of different sequences interspersed across the spike glycoprotein, it becomes imperative that the constructs be evaluated. In this endeavor, we utilized the Expasy ProtParam Tool and based our assumptions on the PEST hypothesis and the Instability Index determinants, which include analysis of the residues that make any biomolecule have lower half-life or increased instability (includes Proline(P), Glutamic acid (E), Serine (S), Threonine (T), Methionine (M) and Glutamine (Q)) in the designed constructs and including Guruprasad’s basis[51] that the relative abundance of Asparagine (N), Lysine (K) and Glycine (G) has been found in stable proteins (**Table 4B**). Moreover, basic or neutral isoelectric points also indicate a stable *in vivo* half-life for the modeled constructs. Since we cannot assess it entirely on these assumptions, we investigated the stability of the constructs through a 100 nanoseconds Molecular Dynamics Simulation. But due to constraints of computational time and effort, we assessed the stability of Construct_4 and 5 since they gave the best estimate of stability based on our assumptions as listed in **Table 4(A, B, C)**. The abovementioned parameters went into this assumption coupled with the Ramachandran Plot, ERRAT-3D, and the Z-score. **Figure 5A** and **Supplementary Figure 3A** indicates the RMSD fluctuation of the stable vaccine constructs 4 and 5, which have been demarcated as the most stable constructs based on these parameters.

**Table 4(A):**
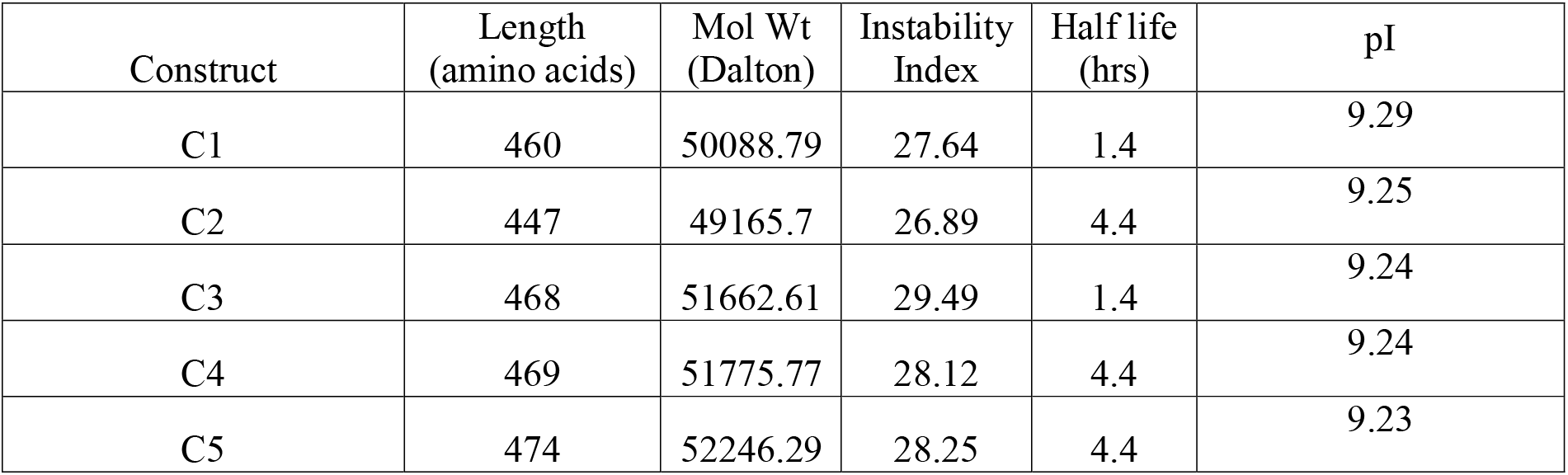
The ProtParam based physicochemical based validation of vaccine constructs

**Table 4(B):** Assessment of Stability based on PEST hypothesis and Guruprasad’s observations.

| Construct | P | E | S | T | Q | M | NKG |
| --- | --- | --- | --- | --- | --- | --- | --- |
| C1 | 6.5 | 5.4 | 4.8 | 9.8 | 3.9 | 0.4 | 19.5 |
| C2 | 7.2 | 5.4 | 4.5 | 10.1 | 3.8 | 0.2 | 19.4 |
| C3 | 7.1 | 5.3 | 4.9 | 9.6 | 3.8 | 0.4 | 19.2 |
| C4 | 7 | 5.3 | 4.9 | 9.6 | 3.8 | 0.4 | 19.2 |
| C5 | 7 | 5.5 | 4.9 | 9.5 | 3.8 | 0.4 | 19.2 |

**Table 4(C):** Validation of Modeled vaccine Constructs

| Construct | Ramachandran Plot |  | Z Score | ERRAT<br>3D | WHATCHECK |
| --- | --- | --- | --- | --- | --- |
|  | Allowed + Favored regions<br>(in percent) | Disallowed<br>regions(in<br>percent) |  |  |  |
| C1 | 91.5+7.3+0.8 | 0.5 | -4.02 | 88.1395 | Pass |
| C2 | 93.2+5.7+0.3 | 0.8 | -4.59 | 83.698 | Pass |
| C3 | 88.9+10.9+0.0 | 0.2 | -4.31 | 86.0045 | Pass |
| C4 | 92.1+6.9+0.2 | 0.7 | -4.62 | 90.4444 | Pass |
| C5 | 93.1+5.1+0.5 | 1.2 | -5.57 | 87.3085 | Pass |

**Figure 5:**
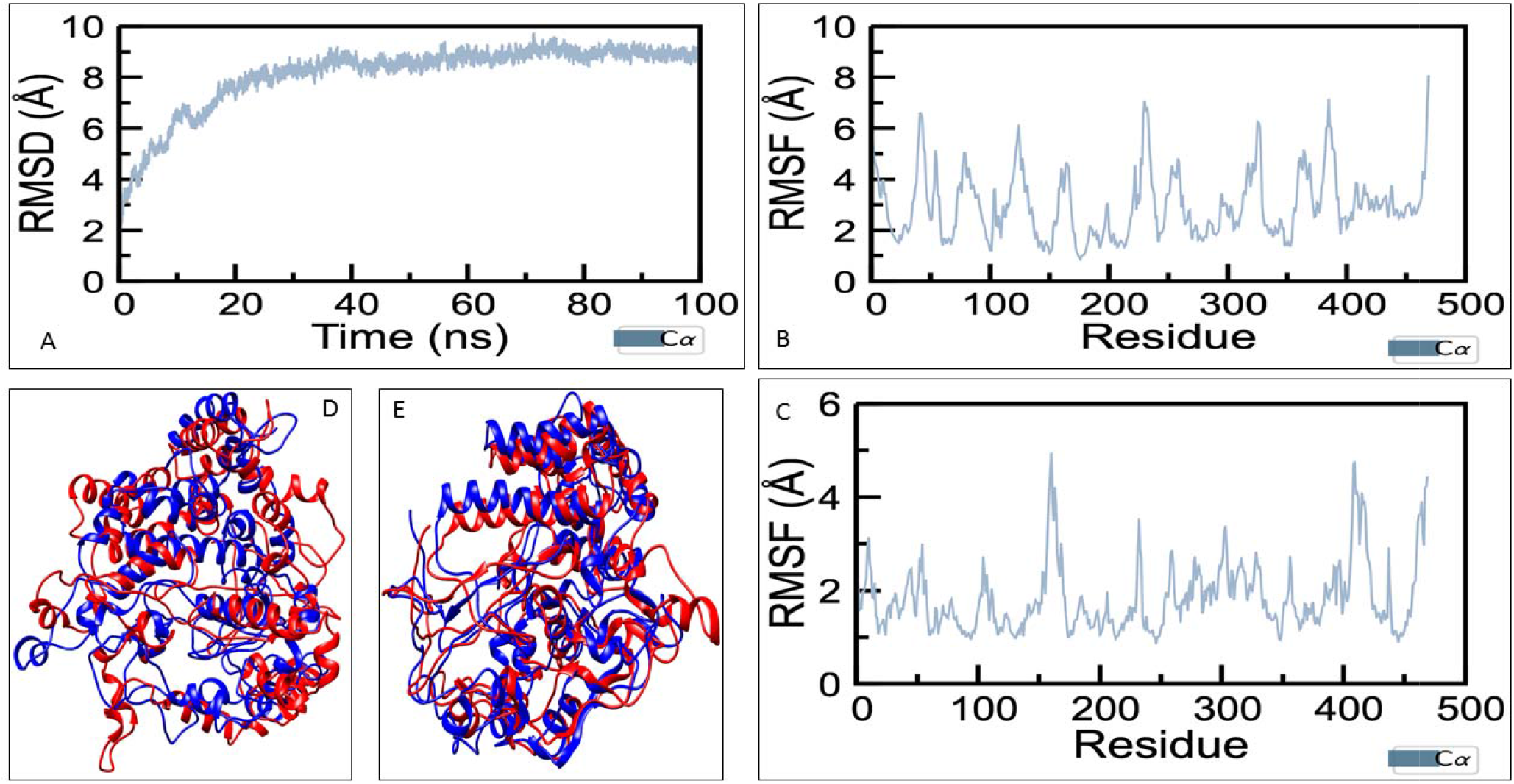
**A**: C-alpha RMSD of Vaccine Construct_4 for a 100ns MD simulation; **B:** RMSF plot comprising C-alpha atoms based on Vaccine Construct_4 from 0-30 ns; **C:** RMSF plot comprising C-alpha atoms based on Vaccine Construct_4 from 30-100 ns, the peaks represent the regions where the loop is in abundance based on residue sequence (across residue index); **D:** Initial rearrangement is depicted through superimposed frames of Vaccine Construct_4 at 0 ns (in red) and at 30 ns (in blue); **E:** Rearrangement through superimposed frames at 30 ns (in red) and at 100 ns (in blue).

### 3.8 MD of vaccine construct

During the initial phase of MD simulation, subsequent conformational changes in the vaccine construct were observed, as illustrated in **Figure 5A**. During the rest of the simulation time average Root Mean Square Deviation (RMSD) of the C-alpha atoms remained constant around 9Å **(Figure 5A)**. This suggests that after the initial structural rearrangements, the vacci**ne** construct achieved a stable three-dimensional structure, as evident in **Figure 5A**, and the superimposed structures obtained at 0 ns and 30 ns in **Figure 5D,** followed by another at 30 ns and 100 ns in **Figure 5E**. Further, to identify the regions of the vaccine construct that contribute to structural rearrangements, Root Mean Square Fluctuations (RMSF) of C-alpha atoms were calculated at 0-30 ns and another from 30-100 ns in **Figures 5B and 5C,** respectively.

The residue fluctuation is due to the low distribution of stable secondary structure rearrangements in the region between residues 153-176 and 412-424, accompanied by an abundance of loops, as observed from the RMSF plots. The abundance of loops confers its affinitive ability for its complementary receptor. An evaluation of the secondary structure features present in the vaccine construct was evaluated through a Dictionary of Secondary Structure Plot (DSSP)[82] in **Supplementary Figure 1**.

The simulation indicates that Construct_4 conforms to the stability directives that are expected of it and can be reproduced under experimental conditions. Considering the MD, we extrapolated our results to include Construct 5 as a suitable vaccine construct that can be considered in the development of vaccines since they scored on a similar index across the parameters on which we assessed and went ahead with the simulation study of Construct_4. MD simulation of vaccine construct 5 was carried out in a similar manner **(Supplementary Figure 3)**.

### 3.9 Docking of the construct with TLR-4

Based on the MD simulation and the observed conformational changes in the modeled vaccine construct 4, molecular docking was carried out between TLR4 receptor and the vaccine construct 4 through Patchdock[66], which carries out docking of rigid molecules based on a protocol that employs molecular shape representation, surface patch mapping, followed by filtering and scoring. A semi-flexible docking protocol was employed involving the vaccine constructs post-simulation and the associated TLR-4 structure. The obtained results were refined and ranked using Firedock[67] based on a global energy value (in kJ/mol) that helps determine the binding affinity of the molecules being considered. The results indicate that the post-simulation Construct_4 does bind TLR-4 with a high binding affinity **(Supplementary Table 2)** with a Global Energy Value (which is an analog of ranking based on binding affinity) of −24.18 kJ/mol. The interface between Construct_4 and TLR4 was analyzed post-docking through UCSF Chimera[12] and DimPlot[83]. These mainly include the non-covalent interactions that are determined through distance constraints between the two docked molecules and involve hydrogen bonds and hydrophobic interactions across the interface. Each of the residues has been specified that contribute to these interactions has been visualized as spheres and depicted on the basis of the corresponding non-covalent interaction across the protein-protein interface in **Figure 6A**. A similar docking was carried out between the modeled vaccine construct 4 (pre-MD simulation) with TLR-4 with a comparable affinity value (−9.10 kJ/mol). The docking with the simulated vaccine construct 4 implicates the following residues E187, K203, E255, Y311, N335, N336, Y414 and K417 involved in H-bonds across the interface with TLR-4. The docking with the initial vaccine construct 4 shows the interface residues involved in interacting with TLR4 includes S330, C371, S372, Y414, G328, S372, L375, Y391 and Y414 among others **(Supplementary Figure 4B)**. This can be attributed to the post-simulation transition that is represented in the structural rearrangement (**Figure 5E**). The change in residues forming hydrogen bonds across the interface indicates the role of polar residues like E187, K203, N335. This change in the residue pattern can conclusively indicate that the docking pose had changed significantly post the 100ns simulation run **(Supplementary Figure 4A and Figure 6A)**. The docking with the modeled vaccine construct 5 indicated residues like T258, G261, P262, G263, P264, G265, A266, I2667, T269 are involved in binding with TLR-4 only **(Supplementary Figure 2)**.

**Figure 6:**
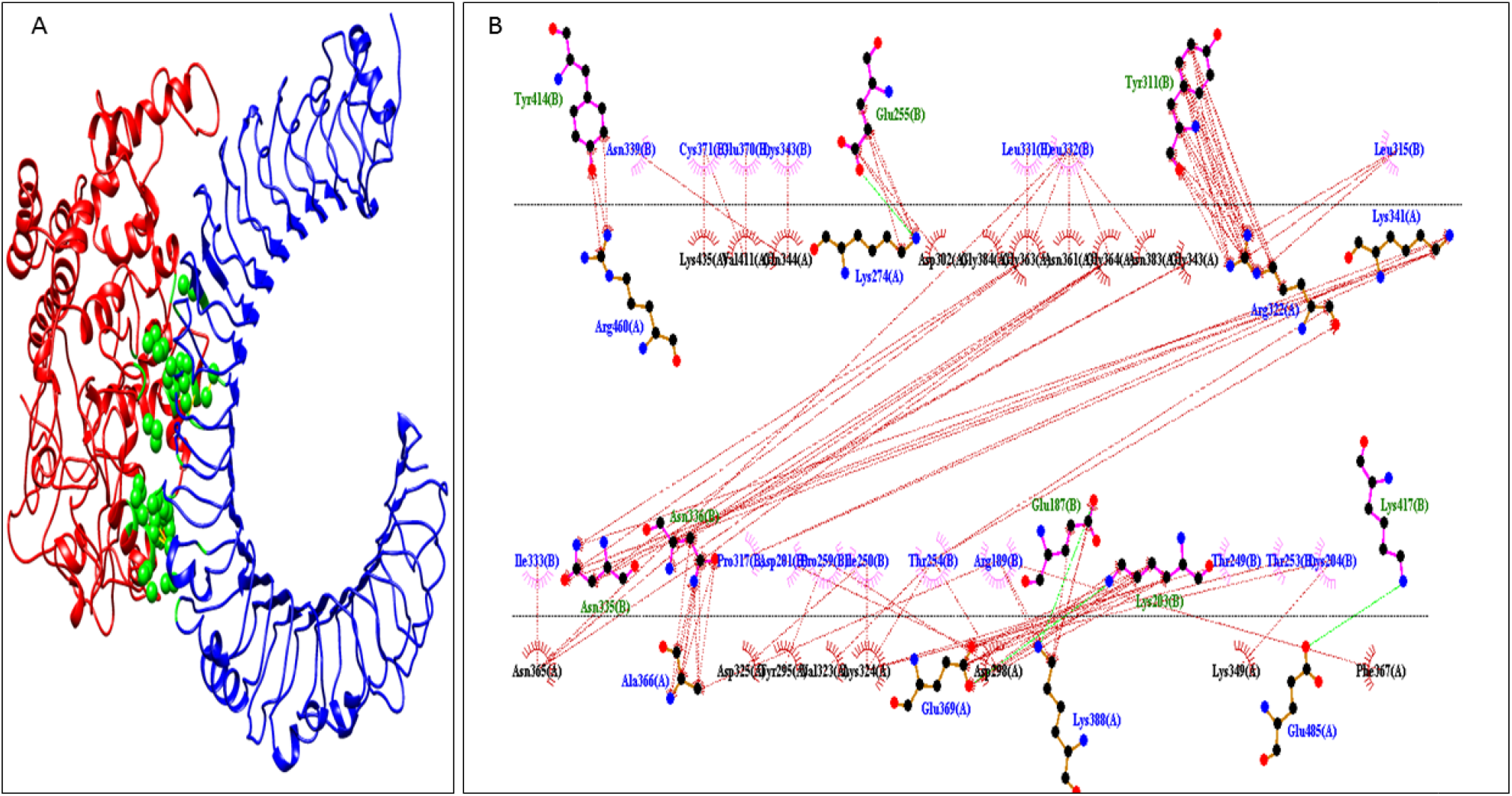
**(A)** The interface of the Vaccine Construct 4 (Chain B: red) and TLR4 (Chain A: blue) are marked by the residues involved across the protein-protein interface through UCSF Chimera. The residues have been marked by green-colored spheres in between;(**B)** Each residue across the interface has been labeled and the corresponding interactions they take part in across the interface through DimPlot.

The docking can be conclusively used to assume that the binding of the vaccine construct to the TLR receptor may elicit signal cascades responsible for the desired immune response towards conferring a suitable prophylactic effect. The utilization of servers that allow simulation of immune response generated based on vaccine dosage was not done in this regard owing to the limitations of the predictions across experimental settings.

### 3.10 MM/GBSA based evaluation of Docked Pose

Molecular Mechanics/Generalized Born Surface Area (MM/GBSA) methods are utilized **to** determine binding free energies for macromolecules and have been used efficiently in t**he** scoring of protein-protein interactions and suitably circumvent the computationally extensi**ve** free-energy calculation methods. Datasets employed to validate their role by comparing th**eir** scoring based on experimentally solved protein-protein interactions indicates the superiority **of** MM/GBSA based calculation models in determining correct docking poses. MM/GBSA h**as** been evaluated to be better than the Molecular Mechanics/Poisson-Boltzmann Surface Area (MM/PBSA) when evaluating protein-protein interactions[68]. The HawkDock Server[68] has been used explicitly in this purpose post-docking to evaluate the docked pose as being accurate. The residues that were implicated across the protein-protein interface on LigPlot of the top-scoring pose by FireDock have been assessed by a scoring function based on force-field parameters influenced through MM/GBSA calculations with similar residues being implicated. The overall binding free energy of the TLR-4 and vaccine construct was predicted to be −23.18 kcal/mol (**Supplementary Table 3**).

## 4. Discussion

One of the most potent options that are being explored to curtail the spread of SARS-CoV-2 includes the design and development of appropriate vaccines. The entire process of vaccine development involves an extensive timeline ranging from experimental to clinical settings. In recent times, with advancement in molecular immunology and the development of several epitope mapping methods, several semi-empirical approaches at vaccine design have been introduced. Multi-epitope-peptide-based subunit vaccines based on predicted B and T cell epitopes utilizing similar bioinformatics tools has made remarkable strides. Despite the cons of low immunogenicity and multiple doses, the trade-off with eliciting neutralizing antibodies, humoral immune response, and relative safety when associated with attenuated or inactivated virus vaccines plays in favor of these approaches[84].

Looking into the probable limitations, strategies have been employed to design an epitope-based peptide vaccine utilizing the major subunits of the spike glycoprotein of SARS-CoV-2. In selecting the concerned immunogen, S1 and the S2 domains of the surface glycoprotein have been utilized separately, keeping in mind the limitations associated with using the entire surface glycoprotein[69, 85]. The prediction of the three different types of epitopes (B cells, T_HTL_ and T_CTL_) by utilizing multiple servers in each case based on different algorithms and selection based on a consensus involving all the servers, including experimental basis behind alleles (HLA Class I and Class II), helps add sustenance to the concluded list of epitopes[70–74]. The predicted epitopes were found to be experimentally identified across a recent microarray study of mapped epitopes on the surface glycoprotein [86]. Each of the predicted epitopes was verified based on described parameters and were found to be antigenic, non-allergenic, nontoxic, and share high coverage across the spike glycoprotein of SARS-CoV-2 sequences[37]. The epitope sequences were also matched with a recently deposited SARS-CoV-2 sequence from Gujarat, India (QJC19491.1) and showed complete sequence coverage for all Indian sequences of the spike glycoprotein deposited to date. Based on the necessity of an immunostimulant, an adjuvant’s role becomes important, and specifically, a chimeric adjuvant comprising TLR-4 agonists was designed and validated to have overlapping B cell epitope regions through the IEDB server. The adjuvant sequences have been experimentally evidenced to generate neutralizing antibodies, cytokines, and natural killer cells. A rigorous assessment of the implicated toll-like receptor in the case of SARS-CoV-2 has been carried out before deciding upon the use of TLR4 agonists which is based mainly on the association of TLR-4 with viral surface glycoprotein in SARS-CoV and Respiratory Syncytial Virus (RSV).The adjuvant and the predicted epitope sequences were appropriately joined by rigid linkers based on their arrangement. Even the chimeric adjuvant sequences were linked by short rigid linkers (XP)_n_ to aid in ensuring stability. GPGPG and AAY linkers were used as intra HTL and CTL linkers, respectively. The separation between the vaccine components has been arrived upon by the role of EAAAK as a linker between adjuvant and the peptide epitope sequences[43, 44]. The designed vaccine constructs based on the arrangement of the chimeric adjuvant were validated and concluded to be highly antigenic, non-allergenic, and nontoxic. This makes up for the limitations of subunit vaccines as mentioned above.

The stability of the vaccine constructs was assessed based on physico-chemical parameters that make it suitable for purification across experimental settings. The presence of cysteine bonds (without introducing any engineered mutations) points to the stability of the modeled constructs. The presence of defined secondary structure characteristics across the modeled construct, mainly strand and loop regions, has led us to verify the dynamic stability through a 100ns MD simulation run. Moreover, the assessment of stability through amino acid composition[51, 52], correlation with in vivo half-life periods, and calculated isoelectric points[53] help us identify three of the five modeled vaccine constructs to exhibit similar stability parameters. The utilization of the ProSA web server[49], SAVES server for Z-score and Ramchandran plot[48], ERRAT-3D[50] scores, helps validate the modeled constructs. Based on the arrangement of the adjuvant, CTL, HTL and B cell epitopes, five different vaccine constructs were considered and modeled as mentioned in Supplementary Tables 1A and 1B. The utilization of linkers interspersed between the different regions allows for a suitable stable vaccine construct, as indicated through Tables 3 and 4. Overlaps based on T cell and B cell epitopes were kept in mind to ensure continuation, as indicated in Tables 1. The rationale behind the vaccine construct is primarily based on the thermo-stability of the designed protein sequence. Rearranging the vaccine constructs based on the different predicted epitopes would lead to a marked decrease in stability of the protein construct as is indicated in Tables 3 and 4. The vaccine construct would be ideally endocytosed and broken up into smaller peptides before presentation through antigen presentation systems that elicits a specific response against the peptides presented. Hence, rearranging the vaccine construct would lead to an unexpected change in generated immune response.

A semi-flexible docking of TLR-4 and Construct_4 was carried out. Although there are no experimental data to insinuate any such interaction but the structural features of the vaccine construct and the functional characteristics of TLR-4, an accepted dock model can be achieved in this scenario. Moreover, the N terminal comprises TLR-4 agonists which helps compound the interaction between the two proteins. Based on TLR-4 PDB structures[63] and their corresponding binding sites followed by Ellipro[21] and Castp[65] predictions of binding sites over the receptor, docking was carried out with the vaccine construct over PatchDock[66] server, and the top pose based on binding affinity values was selected. The interacting residues at the protein-protein interface were mapped. The interaction occurs over the surfaces of TLR-4 which has a high electrostatic potential. The mapped non-covalent interactions are sufficient to indicate probable binding to the receptor surface through DimPlot and visualization through UCSF Chimera. The post docking MM/GBSA evaluation validates the docked pose and the involved residues across the protein-protein interaction interface.

The binding of B cell epitope sequences in the vaccine construct with the predicted B cell regions over TLR-4 will help in the generation of innate immune responses through plasma B cells producing antibodies and memory B cells that confer long-lasting prophylactic response against the virus. Moreover, recent MD studies have indicated that the trimer of spike protein is more stable as compared to the monomer, and hence may influence the stability of spike-based vaccines.

Since no experimental data is evidenced to verify the docking of the TLR4 and the vaccine construct, a theoretical determination of the protein-protein interaction between them may be carried out by running extensive MD simulations between both the proteins (tempered binding) and determining their association and dissociation profiles alongside residence time to assess the best docking pose of the two proteins in their energy minimized conditions. This may circumvent the limitations associated with the unavailability of experimental evidence to a certain extent, which could not be carried out due to computational constraints.

## 6. Authors’ contributions

SS and DM conceived the idea of this study and designed the experiments together. DM performed the experiments. AJ, SS, DM and JP analyzed the data. DM, SS, AJ and JP contributed to drafting the manuscript.

## 7. Acknowledgement

AJ acknowledges the Department of Biotechnology, Govt of India for the Ramalingamswami Re-entry Fellowship-2019.

## Supplementary Data

**Supplementary Table 1A:**
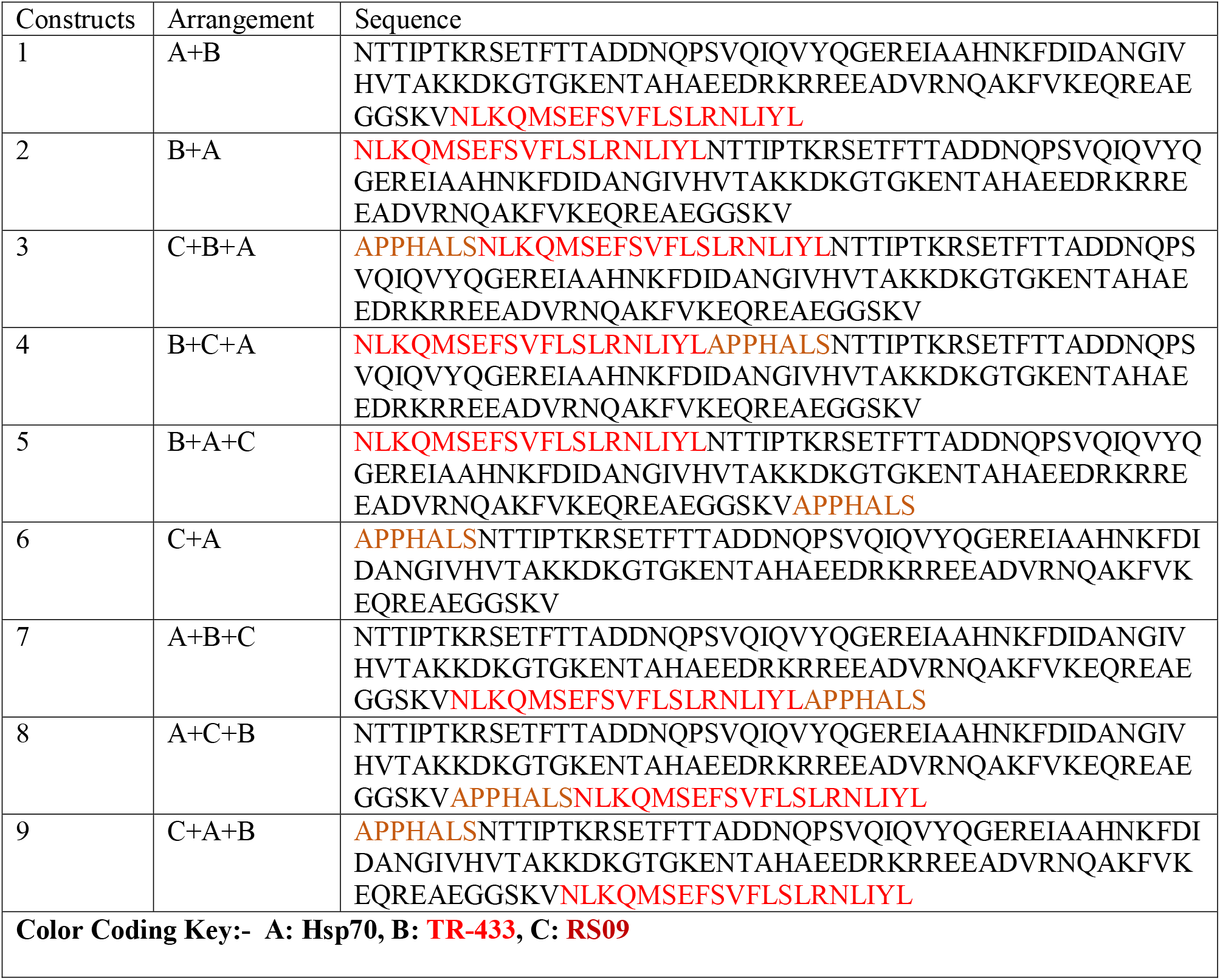
Initial adjuvant arrangements considered

**Supplementary Table 1B:**
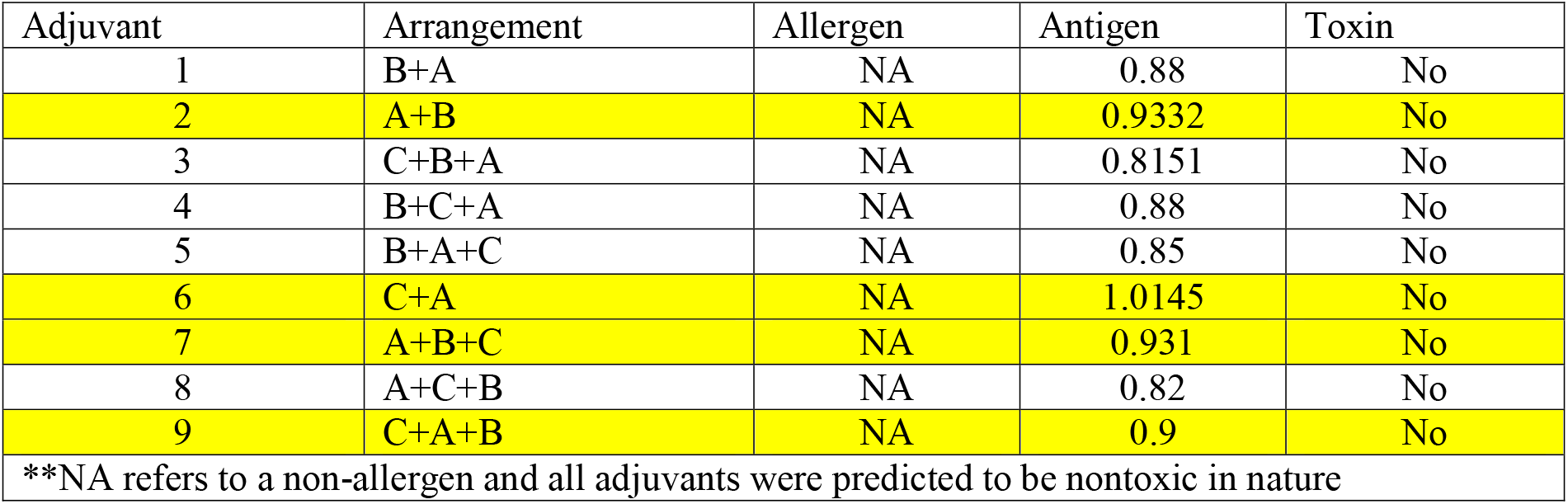
The 2^nd^, 6^th^, 7^th^ and 9^th^ construct were found to be the most antigenic (a stringent Vaxijen score of =>0.9 whereas threshold lies at 0.4) and were utilized to design the vaccine constructs

**Supplementary Table 2:**
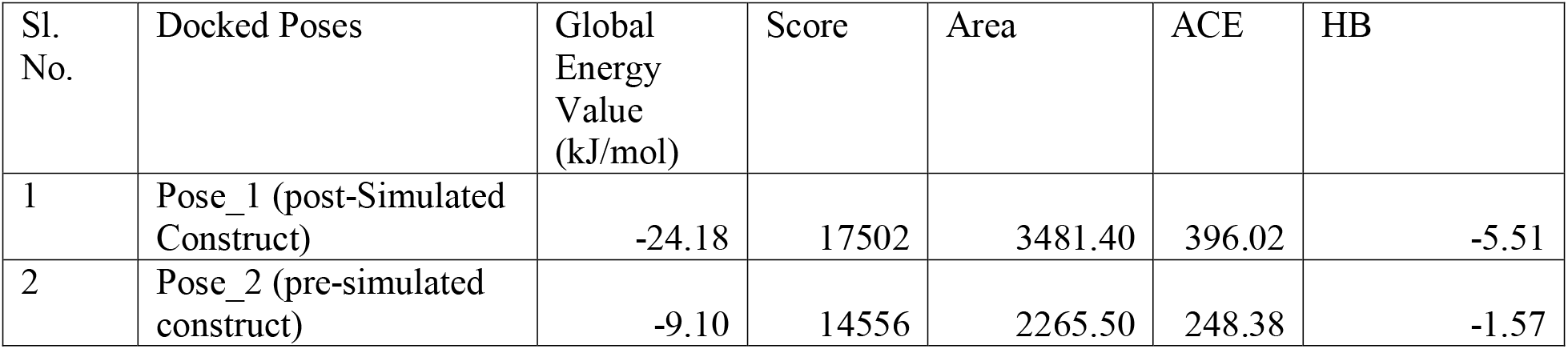
Patchdock mediated docking between Construct_4 and TLR-4 with Ranking by FireDock based on Global Energy Value.

**Supplementary Table 3:**
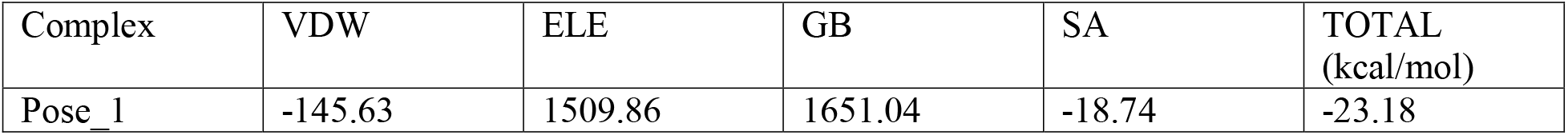
MM/GBSA of the docked pose of the TLR-4 and the vaccine construct 4 has been evaluated across Van der Waal potentials (**VDW**), electrostatic potentials (**ELE**), polar solvation free energies predicted by the Generalized Born model (**GB**) and the non-polar contribution to the solvation free energy calculated through an empirical model (**SA**).

**Supplementary Figure 1:**
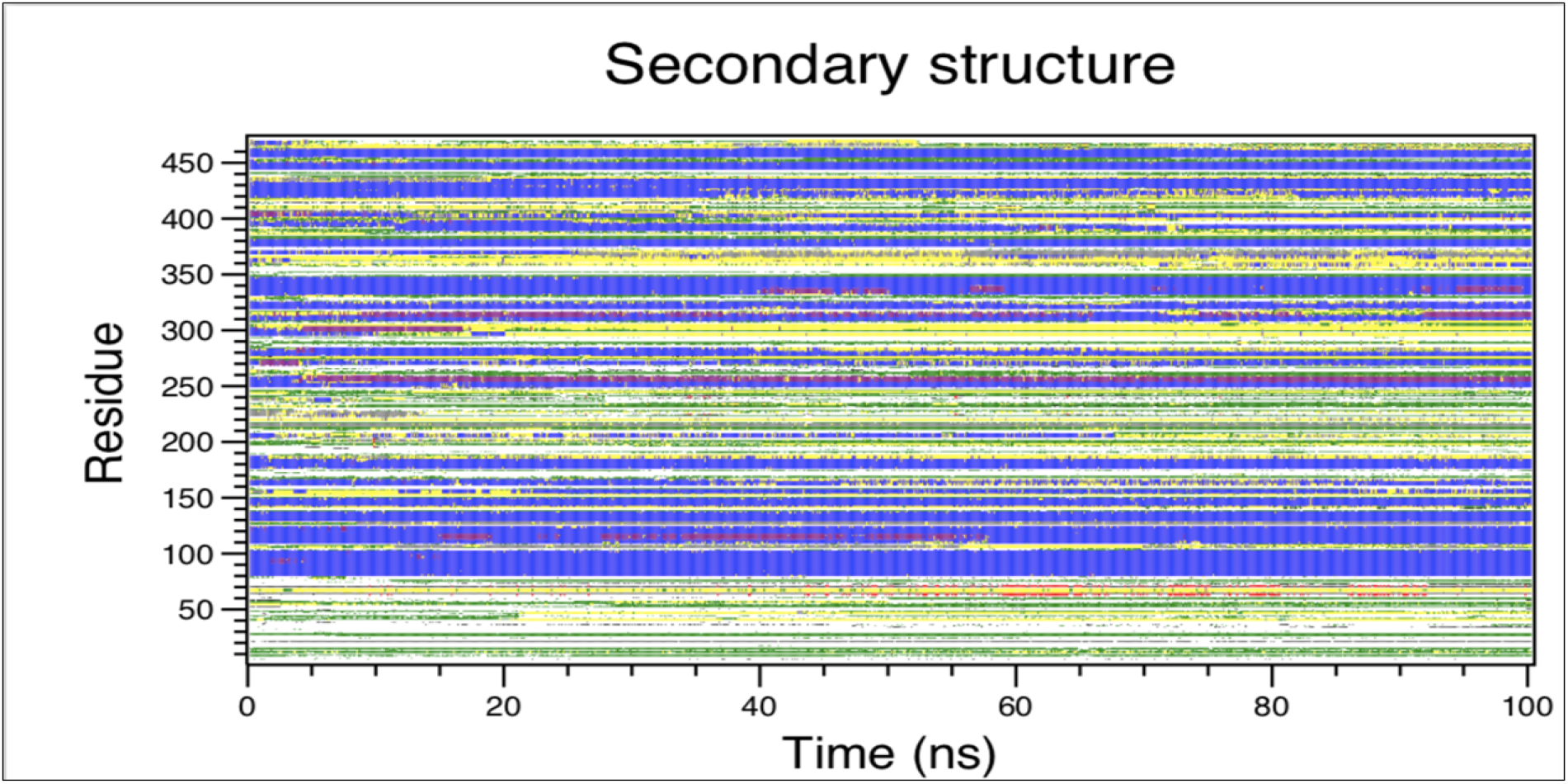
The DSSP plot of Vaccine Construct_4 from 0-100ns

**Supplementary Figure 2:**
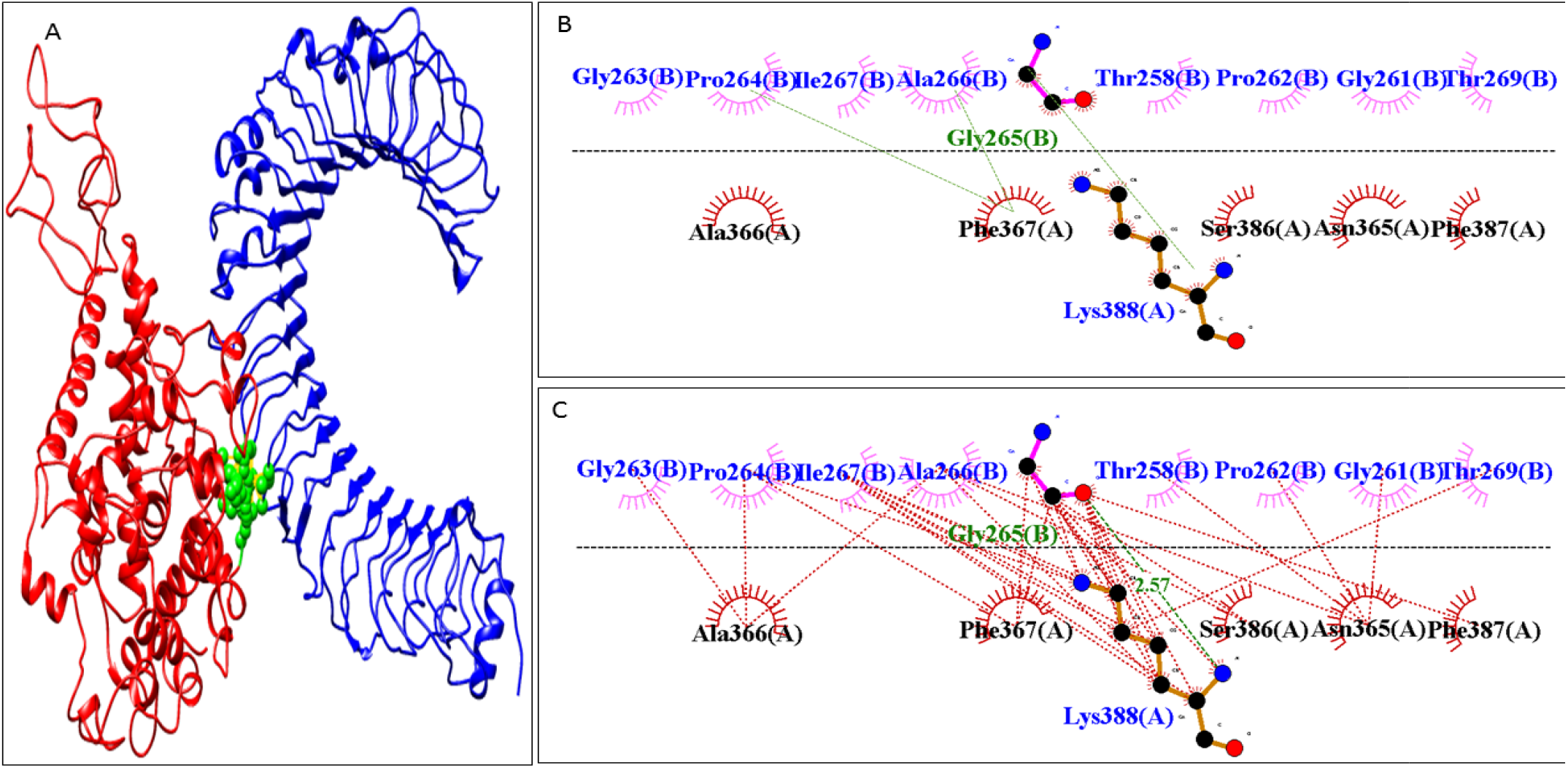
The residues which contribute to interactions across the docked protein-protei: interface of Vaccine Construct 5 and TLR4 **A:** Vaccine Construct 5 (**red**) and TLR4 (**blue**).Green sphere mark the interface between the TLR4 and Vaccine Construct 5 which contribute to the protein-protein interaction; **B:** Hydrogen bonds between the TLR-4 and the Construct 5 are marked by green lines; C Hydrophobic interactions between the TLR-4 and the Construct 5 which contribute to the protein-protein interaction from DimPlot.

**Supplementary Figure 3:**
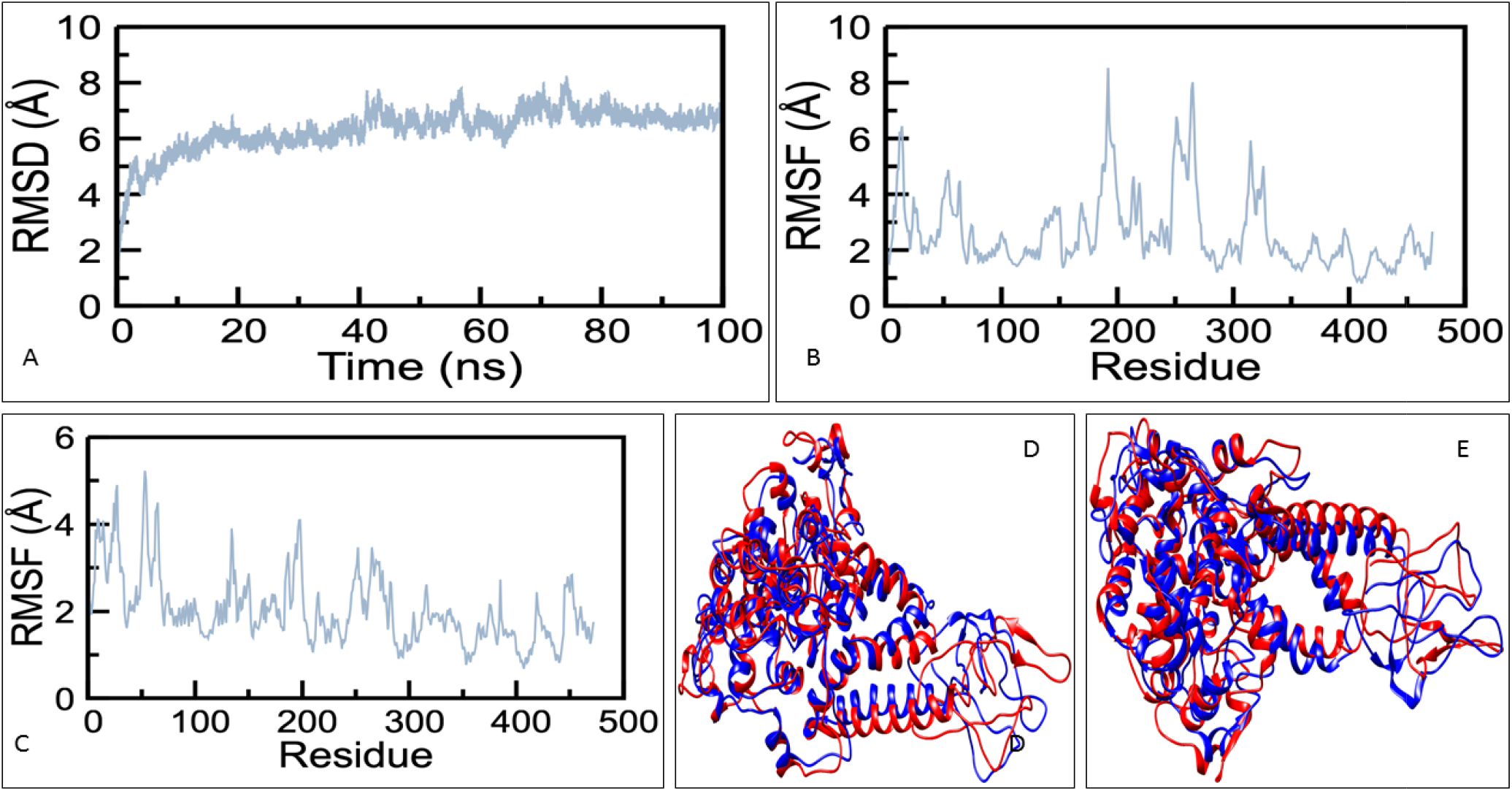
(**A**) C-alpha RMSD of Vaccine Construct_5 for a 100ns MD simulation; **(B):** RMSF plot comprising of C-alpha atoms based on Vaccine Construct_5 from 0-30 ns; **(C)** RMSF plot comprising of C-alpha atoms based on Vaccine Construct_5 from 30-100 ns, the peaks represents the regions where loop is in abundance based on residue sequence (across residue index); **(D)** Initial rearrangement is depicted through superimposed frames of Vaccine Construct_5 at 0 ns (in red) and at 30 ns (in blue); **(E)** Rearrangement through superimposed frames at 30 ns (in red) and at 100 ns (in blue).

**Supplementary Figure 4:**
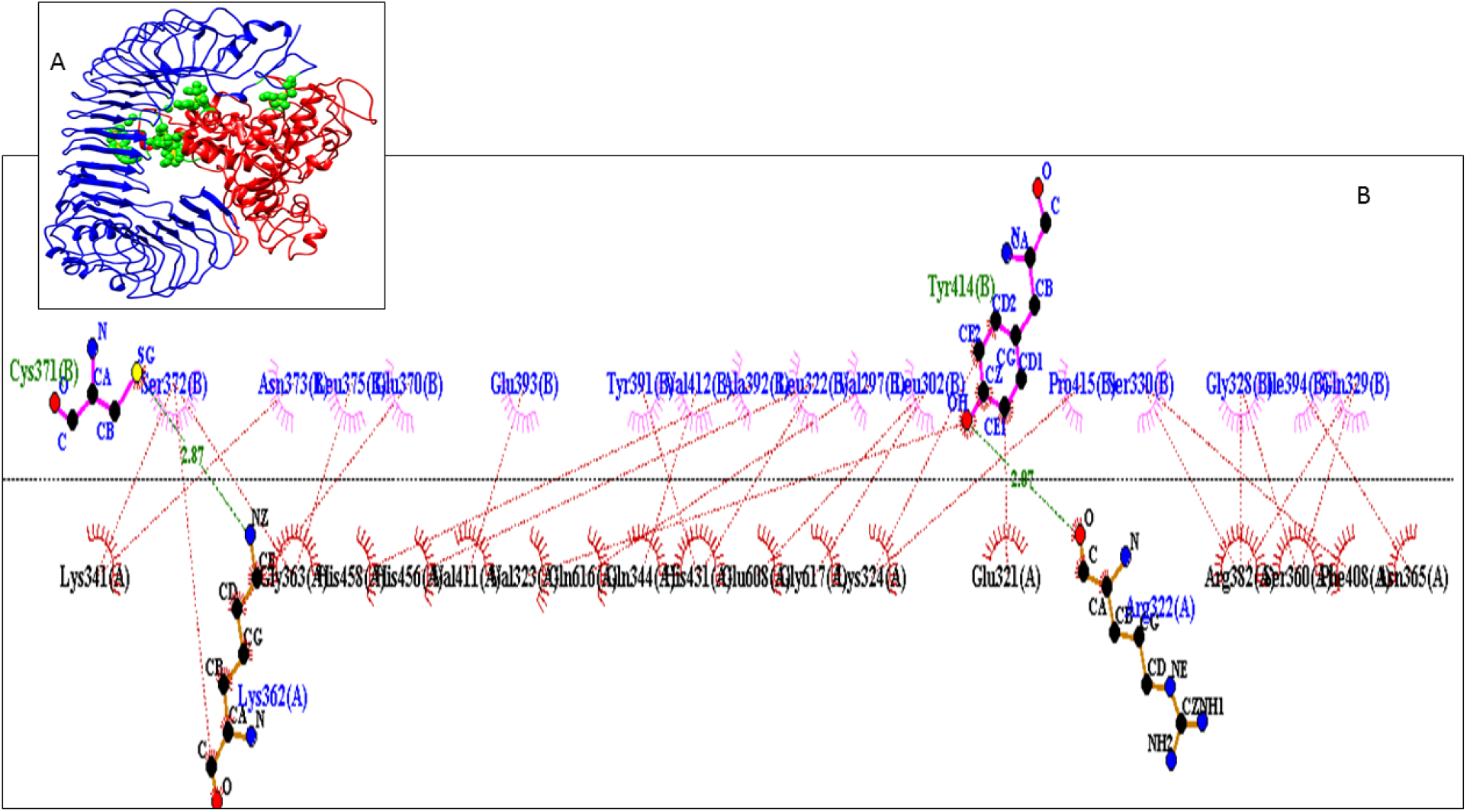
The docked poses generated from: (A) Docking between TLR-4 (blue) and designed vaccine construct 4 (red); (B) Residues involved in interaction between the TLR4 and the designed vaccine construct 4.

