## Supplementary Information for "*In silico* design of Multi-epitope-based peptide vaccine against SARS-CoV-2 using its spike protein"

**Supplementary Data**

**Supplementary Table 1A:** Initial adjuvant arrangements considered

| Constructs | Arrangement | Sequence |
| --- | --- | --- |
| 1 | A+B | NTTIPTKRSETFTTADDNQPSVQIQVYQGEREIAAHNKFDIDANGIVHVTAKKDKGTGKENTAHAEEDRKRREEADVRNQAKFVKEQREAEGGSKVNLKQMSEFSVFLSLRNLIYL |
| 2 | B+A | NLKQMSEFSVFLSLRNLIYLNTTIPTKRSETFTTADDNQPSVQIQVYQGEREIAAHNKFDIDANGIVHVTAKKDKGTGKENTAHAEEDRKRREEADVRNQAKFVKEQREAEGGSKV |
| 3 | C+B+A | APPHALSNLKQMSEFSVFLSLRNLIYLNTTIPTKRSETFTTADDNQPSVQIQVYQGEREIAAHNKFDIDANGIVHVTAKKDKGTGKENTAHAEEDRKRREEADVRNQAKFVKEQREAEGGSKV |
| 4 | B+C+A | NLKQMSEFSVFLSLRNLIYLAPPHALSNTTIPTKRSETFTTADDNQPSVQIQVYQGEREIAAHNKFDIDANGIVHVTAKKDKGTGKENTAHAEEDRKRREEADVRNQAKFVKEQREAEGGSKV |
| 5 | B+A+C | NLKQMSEFSVFLSLRNLIYLNTTIPTKRSETFTTADDNQPSVQIQVYQGEREIAAHNKFDIDANGIVHVTAKKDKGTGKENTAHAEEDRKRREEADVRNQAKFVKEQREAEGGSKVAPPHALS |
| 6 | C+A | APPHALSNTTIPTKRSETFTTADDNQPSVQIQVYQGEREIAAHNKFDIDANGIVHVTAKKDKGTGKENTAHAEEDRKRREEADVRNQAKFVKEQREAEGGSKV |
| 7 | A+B+C | NTTIPTKRSETFTTADDNQPSVQIQVYQGEREIAAHNKFDIDANGIVHVTAKKDKGTGKENTAHAEEDRKRREEADVRNQAKFVKEQREAEGGSKVNLKQMSEFSVFLSLRNLIYLAPPHALS |
| 8 | A+C+B | NTTIPTKRSETFTTADDNQPSVQIQVYQGEREIAAHNKFDIDANGIVHVTAKKDKGTGKENTAHAEEDRKRREEADVRNQAKFVKEQREAEGGSKVAPPHALSNLKQMSEFSVFLSLRNLIYL |
| 9 | C+A+B | APPHALSNTTIPTKRSETFTTADDNQPSVQIQVYQGEREIAAHNKFDIDANGIVHVTAKKDKGTGKENTAHAEEDRKRREEADVRNQAKFVKEQREAEGGSKVNLKQMSEFSVFLSLRNLIYL |
| **Color Coding Key:- A: Hsp70, B: TR-433, C: RS09** | | |

**Supplementary Table 1B** : The 2^nd^, 6^th^, 7^th^ and 9^th^ construct were found to be the most antigenic ( a stringent Vaxijen score of =>0.9 whereas threshold lies at 0.4) and were utilized to design the vaccine constructs

| Adjuvant | Arrangement | Allergen | Antigen | Toxin |
| --- | --- | --- | --- | --- |
| 1 | B+A | NA | 0.88 | No |
| 2 | A+B | NA | 0.9332 | No |
| 3 | C+B+A | NA | 0.8151 | No |
| 4 | B+C+A | NA | 0.88 | No |
| 5 | B+A+C | NA | 0.85 | No |
| 6 | C+A | NA | 1.0145 | No |
| 7 | A+B+C | NA | 0.931 | No |
| 8 | A+C+B | NA | 0.82 | No |
| 9 | C+A+B | NA | 0.9 | No |
| **NA refers to a non-allergen and all adjuvants were predicted to be nontoxic in nature | | | | |

**Supplementary Table 2**: Patchdock mediated docking between Construct_4 and TLR-4 with Ranking by FireDock based on Global Energy Value.

| Sl. No. | Docked Poses | Global Energy Value (kJ/mol) | Score | Area | ACE | HB |
| --- | --- | --- | --- | --- | --- | --- |
| 1 | Pose_1 (post-Simulated Construct) | -24.18 | 17502 | 3481.40 | 396.02 | -5.51 |
| 2 | Pose_2 (pre-simulated construct) | -9.10 | 14556 | 2265.50 | 248.38 | -1.57 |

**Supplementary Table 3**: MM/GBSA of the docked pose of the TLR-4 and the vaccine construct 4 has been evaluated across Van der Waal potentials (**VDW**), electrostatic potentials (**ELE**), polar solvation free energies predicted by the Generalized Born model (**GB**) and the non-polar contribution to the solvation free energy calculated through an empirical model (**SA**).

| Complex | VDW | ELE | GB | SA | TOTAL (kcal/mol) |
| --- | --- | --- | --- | --- | --- |
| Pose_1 | -145.63 | 1509.86 | 1651.04 | -18.74 | -23.18 |

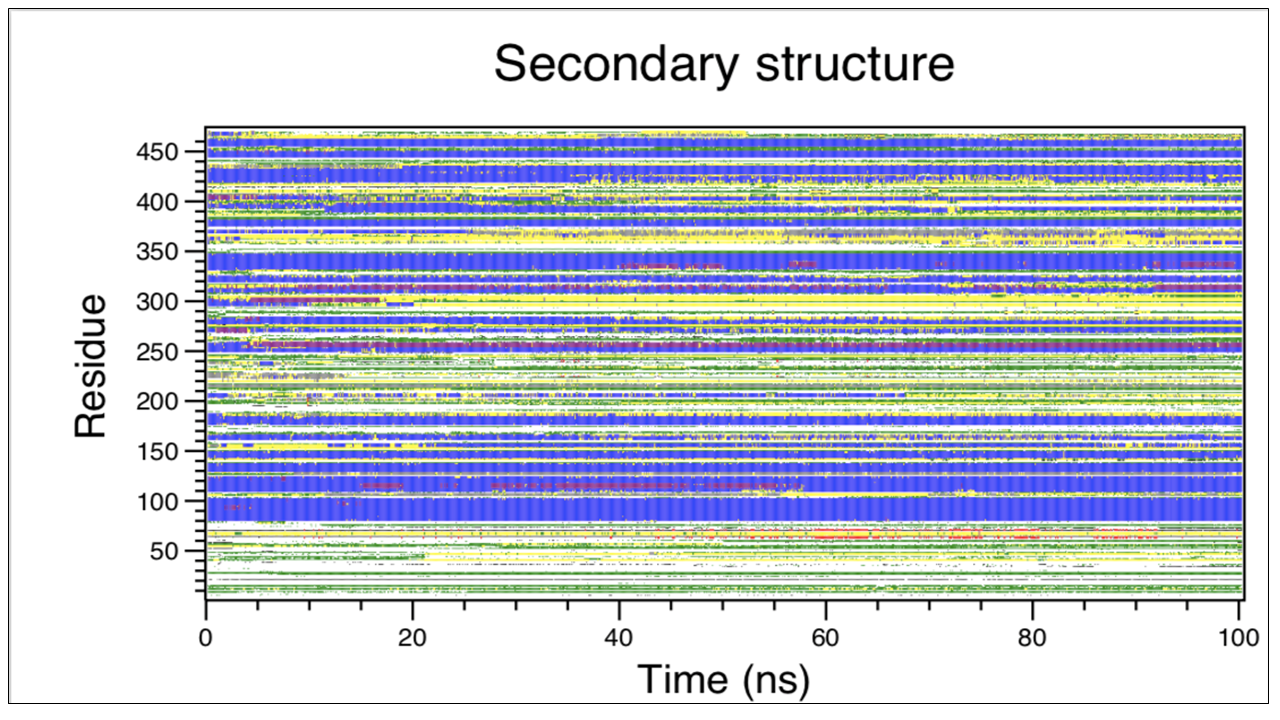

**Supplementary Figure 1:** The DSSP plot of Vaccine Construct_4 from 0-100ns

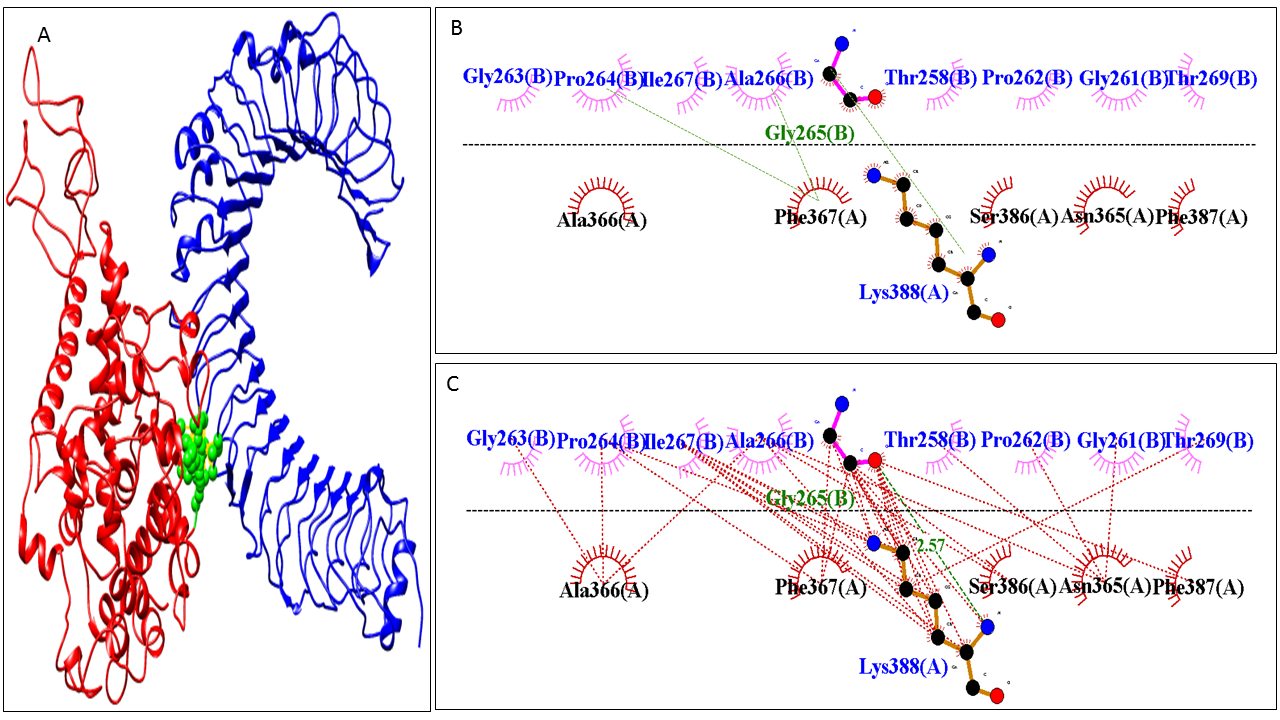

**Supplementary Figure 2:** The residues which contribute to interactions across the docked protein-protein interface of Vaccine Construct 5 and TLR4 **A:** Vaccine Construct 5 (**red**) and TLR4 (**blue**).Green spheres mark the interface between the TLR4 and Vaccine Construct 5 which contribute to the protein-protein interaction; **B:** Hydrogen bonds between the TLR-4 and the Construct 5 are marked by green lines; C: Hydrophobic interactions between the TLR-4 and the Construct 5 which contribute to the protein-protein interaction from DimPlot .

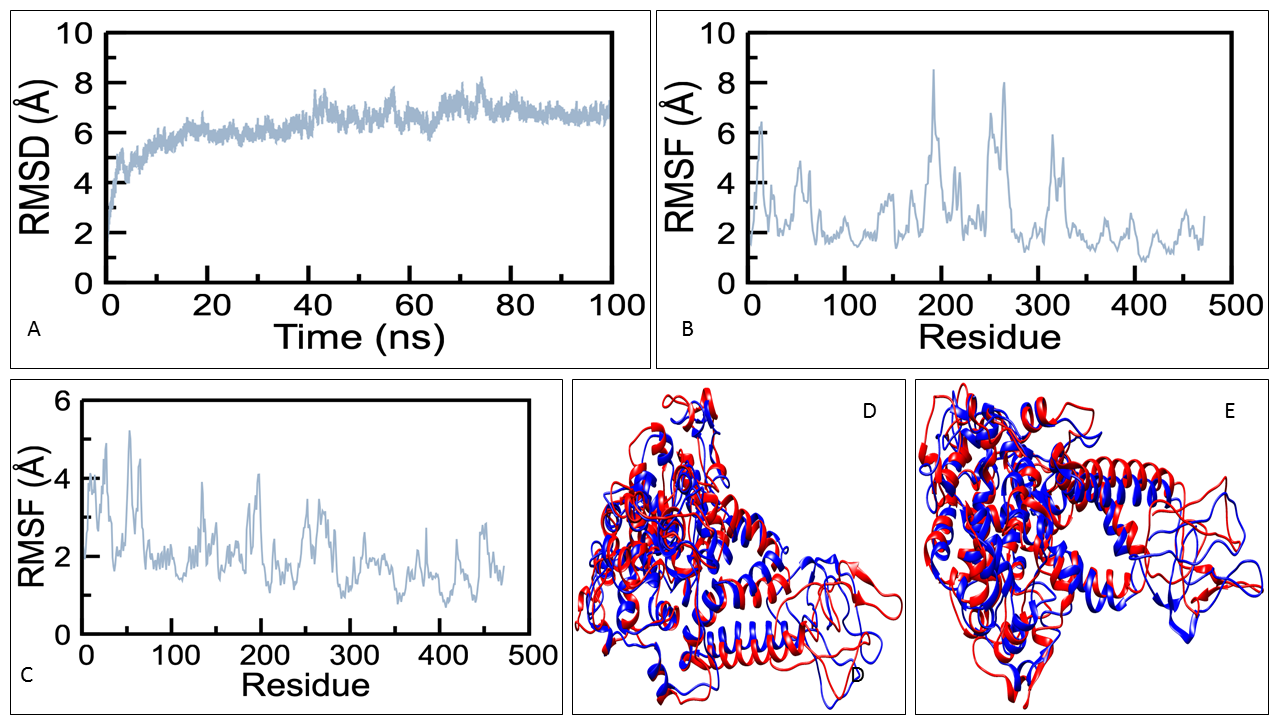

**Supplementary Figure 3:** (**A**) C-alpha RMSD of Vaccine Construct_5 for a 100ns MD simulation; **(B):** RMSF plot comprising of C-alpha atoms based on Vaccine Construct_5 from 0-30 ns; **(C)** RMSF plot comprising of C-alpha atoms based on Vaccine Construct_5 from 30-100 ns, the peaks represents the regions where loop is in abundance based on residue sequence (across residue index); **(D)** Initial rearrangement is depicted through superimposed frames of Vaccine Construct_5 at 0 ns (in red) and at 30 ns (in blue); **(E)** Rearrangement through superimposed frames at 30 ns (in red) and at 100 ns (in blue).

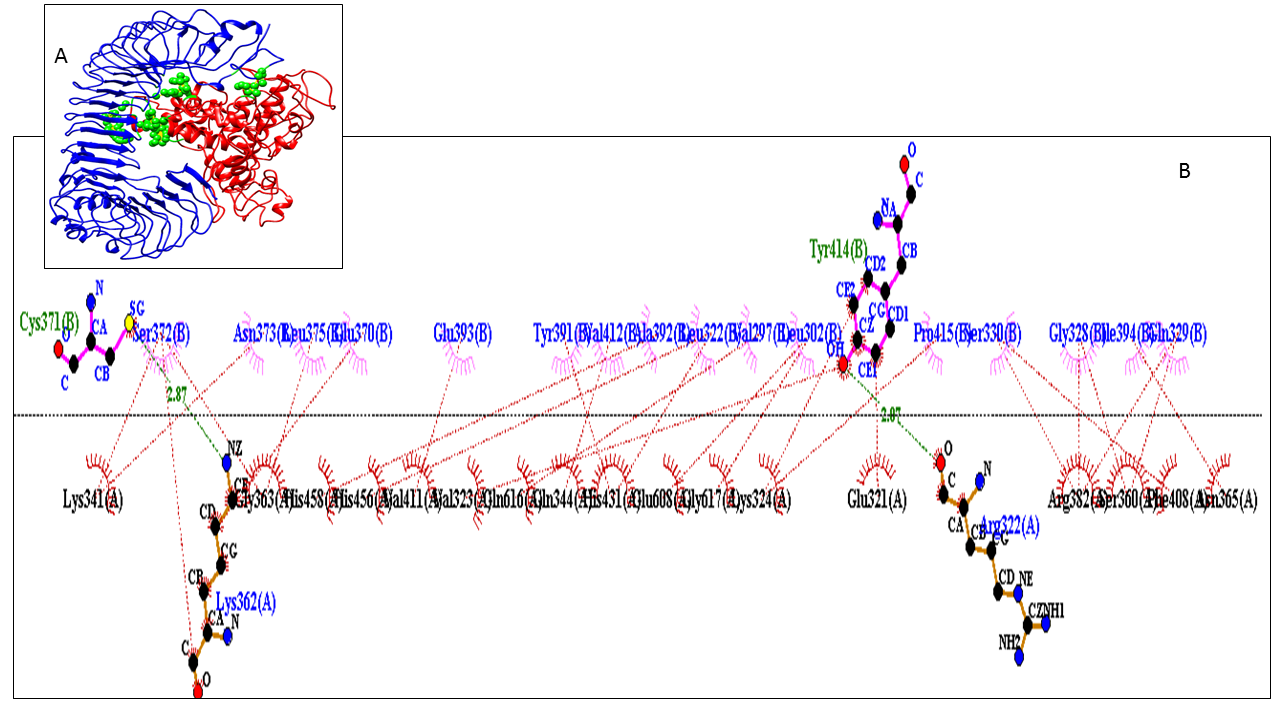

**Supplementary Figure 4:** The docked poses generated from: (A) Docking between TLR-4 (blue) and designed vaccine construct 4 (red); (B) Residues involved in interaction between the TLR4 and the designed vaccine construct 4.
